# Unraveling the influences of sequence and position on yeast uORF activity using massively parallel reporter systems and machine learning

**DOI:** 10.1101/2021.04.16.440232

**Authors:** Gemma May, Christina Akirtava, Matthew Agar-Johnson, Jelena Micic, John Woolford, Joel McManus

**Affiliations:** Department of Biological Sciences; Computational Biology Department Carnegie Mellon University, Pittsburgh PA, USA

## Abstract

Upstream open reading frames (uORFs) are potent *cis*-acting regulators of mRNA translation and nonsense-mediated decay (NMD). While both AUG- and non-AUG initiated uORFs are ubiquitous in ribosome profiling studies, few uORFs have been experimentally tested. Consequently, the relative influences of sequence, structural, and positional features on uORF activity have not been determined. We quantified thousands of yeast uORFs using massively parallel reporter assays in wildtype and Δ*upf1* yeast. While nearly all AUG uORFs were robust repressors, most non-AUG uORFs had relatively weak impacts on expression. Machine learning regression modeling revealed that uORF functions are strongly impacted by both their sequences and locations within transcript leaders. Indeed, alternative transcription start sites highly influenced uORF activity. These results define the scope of natural uORF activity, identify features associated with translational repression and NMD, and suggest that the locations of uORFs in transcript leaders are nearly as important as uORF sequences.

## Introduction

The synthesis of cellular proteins by mRNA translation is an essential process regulated by multiple interactions between *cis-*acting sequences and *trans*-acting factors. Translation initiation is highly regulated to control the rate of protein synthesis. During canonical 5’ cap-dependent initiation, pre-initiation complexes (PICs) assemble at mRNA 5’ ends and scan directionally in search of start codons (Hinnebusch et al., 2016). Due to this directional scanning, mRNA sequences and structures in 5’ transcript leaders have a direct impact on initiation frequency. In particular, upstream Open Reading Frames (uORFs), short coding sequences between the 5’ cap and primary protein coding sequence (CDS), generally decrease the frequency of initiation at downstream CDSs (Wethmar, 2014). Termination at uORF stop codons can also induce nonsense mediated decay (NMD), a major mRNA turnover pathway. As a result, most uORFs are expected to reduce gene expression. Despite this general view, the extent of uORF repression can vary greatly and the underlying causes for this remain unclear.

Despite their general repressive nature, some uORFs enhance expression of their downstream CDS, especially in response to stress (Young and Wek, 2016). A classic example of a gene harboring enhancer uORFs is yeast GCN4, whose transcript leader harbors four uORFs (Mueller and Hinnebusch, 1986). After translation of uORF1, the small subunit of the ribosome remains attached to *GCN4* mRNA, reforming a PIC that subsequently resumes scanning. In the absence of stress, the resulting PIC is rapidly recharged with tRNA-Met in a ternary complex with eIF2 and GTP, which leads to translation of uORFs 2-4 and a corresponding inhibition of *GCN4* CDS translation. However, most stress conditions cause phosphorylation of the eIF2α subunit, which allows rescanning PICs to scan past uORFs 2-4 to translate the *GCN4* CDS (Hinnebusch, 2005). A similar multi-uORF reinitiation system allows stress-dependent translation of mammalian ATF4, in which resumption of scanning after translating a short uORF allows subsequent leaky scanning past a more repressive uORF (Vattem and Wek, 2004). In theory, the same mechanism could allow short uORFs to insulate against longer, more repressive uORFs even in the absence of stress (Lin et al., 2019). However, the frequency of such stress-independent uORF enhancers remains unknown.

Although once thought to be relatively rare, genomic studies have revealed that AUG-initiated uORFs are common, affecting ∼15% of yeast (Ingolia et al., 2009; McManus et al., 2014; Zhang and Dietrich, 2005a) and ∼50% of human genes(McGillivray et al., 2018; Wethmar et al., 2014). In addition to these canonical elements, ribosome profiling studies using drugs that stall initiating ribosomes, have identified even more uORFs that initiate at non-AUG codons (Brar et al., 2011; Ingolia et al., 2011; Spealman et al., 2017). However, other work suggests the drugs used in those studies may exaggerate the frequency of uORF usage (Gerashchenko and Gladyshev, 2014; Kearse et al., 2019; Lareau et al., 2014). Despite the large number of predicted AUG and non-AUG uORFs, most studies focus solely on identifying these elements and few have been experimentally tested (Wethmar et al., 2014). The few non-AUG uORFs that have been tested had relatively modest influences on expression (Spealman et al., 2017; Fan Zhang and Hinnebusch, 2011). Thus, the relative influence of AUG- and non-AUG uORFs on mRNA translation has not been systematically evaluated.

Due to their frequency and functional importance, uORF evolution has been the subject of multiple studies (Zhang et al., 2019). Early work identified 38 genes with AUG-initiated uORFs that were deeply conserved among yeast species by stringent comparisons of draft genome sequences (Zhang and Dietrich, 2005a). Functional evaluations of nine *S. cerevisiae* uORFs showed six altered the expression in a luciferase reporter assay. More recently, genome-wide studies using ribosome profiling identified thousands of AUG and non-AUG uORFs in multiple *Saccharomyces* species, some of which were translated in multiple species (Spealman et al., 2017). Ribosome profiling was also used to identify uORFs used throughout *Drosophila* development, many of which appear to be adaptive due to signs of positive selection detected by comparative genomics (Zhang et al., 2018). Additional analyses of human, mouse, and zebrafish ribosome profiling data estimated uORF regulatory activity using the ratio (U/C) of ribosome footprints in 5’ transcript leaders (UTRs) to footprints in coding sequences (CDS) (Chew et al., 2016; Johnstone et al., 2016). These studies found modest, though statistically significant correlations of U/C ratios in data from mouse and human tissue culture cells and bulk brain tissue, suggesting that uORF activities are somewhat conserved in vertebrates. However, to our knowledge, the regulatory functions of homologous uORFs from different species have not been experimentally compared. Thus, the extent to which uORF functions are quantitatively conserved across species remains unclear.

Currently, the magnitude of uORF activity is thought to depend primarily on the extent to which uORF start codons match the Kozak consensus sequence (Cuperus et al., 2017; Dvir et al., 2013; Noderer et al., 2014; Sample et al., 2019). Analyses of ribosome profiling data from vertebrates found that the U/C ratio was higher for uORFs with start codons in strong Kozak contexts (Chew et al., 2016; Johnstone et al., 2016). Massively parallel reporter assays (MPRAs) of random transcript leaders confirmed that AUG-uORFs in strong Kozak contexts are repressive (Cuperus et al., 2017; Dvir et al., 2013; Ferreira et al., 2014; Noderer et al., 2014; Sample et al., 2019). However, other features have also been shown to affect uORF activity. For example, a uORF-specific MPRA (FACS-uORF) study found rare codons and dicodons within a uORF can determine whether the uORF enhances or represses expression of the CDS (Lin et al., 2019). The structural accessibility of start codons was also found to play a key role in determining the impact of uORFs from the human α-1-antitrypsin mRNA leader (Corley et al., 2017). Other work has shown that uORF activity can depend on the location of uORFs relative to main protein-coding ORF start codons (Beznosková et al., 2013; Grant et al., 2012). However, the relative influences of these features on natural uORF activities have not been determined, primarily because few such uORFs have been functionally assayed.

Similarly, many questions remain regarding uORFs and NMD. The extent to which NMD contributes to typical uORF-mediated gene repression remain unclear, as do the roles of uORF features in determining NMD susceptibility. While uORFs are clearly correlated with NMD (Celik et al., 2017; Smith et al., 2014), the yeast GCN4 and YAP1 uORFs appear to resist NMD (Ruiz-Echevarría and Peltz, 2000). Early studies suggested yeast NMD requires AU-rich *cis*-acting downstream sequence elements bound by Hrp1p (Culbertson and Leeds, 2003; Czaplinski et al., 1999; González et al., 2000; Peltz et al., 1993; Ruiz-Echevarría et al., 1998), however such elements are not well defined and appear to be missing from many NMD targets (Meaux et al., 2008). Other work found premature termination codons (PTCs) were less likely to induce NMD when inserted closer to the 5’ end of a yeast PGK1 mRNA (Muhlrad and Parker, 1999). NMD induction was also shown to be greatly reduced by low-frequency stop codon readthrough at PTCs (Keeling et al., 2004), however high-frequency readthrough did not prevent NMD in another context (Gorgoni et al., 2019). Other studies showed mRNA can be protected from NMD by positioning poly(A)-binding protein close to the termination codon (Amrani et al., 2004; Eberle et al., 2008). Additional work in human cells found translation initiation factors inhibit NMD, as does reinitiation after termination at PTCs (Lindeboom et al., 2016; Raimondeau et al., 2018; Zhang and Maquat, 1997). While these studies helped explain NMD induction by PTCs in coding genes, most did not consider uORFs, which may behave differently due to their extreme 5’ locations in transcript leaders and generally short lengths. Thus, the features that influence uORF induction of NMD remain unclear. Furthermore, the relative importance of NMD and translation inhibition in natural uORF activity has not been systematically evaluated.

Here, we used two MPRA systems, FACS-uORF and PoLib-seq, to quantify the impact of thousands of natural yeast AUG- and non-AUG uORFs on protein expression and ribosome loading. Our results show that most non-AUG uORFs have small impacts on expression compared to AUG-uORFs. Leveraging the massive scale of our results, we evaluated the influence of sequence and positional features of natural AUG-uORFs on their gene regulatory effects. Using a strain deleted of the NMD factor *UPF1* (Δ*upf1*), we showed NMD accounts for roughly a third of uORF repression in yeast, and further investigated how uORF features impact the propensity for NMD. Finally, we used elastic-net regression modeling to select and weight features that have the most significant influences on uORF activity, revealing that uORF location within transcripts leaders plays an important role in determining uORF function. Surprisingly, we found uORF activity often depend on the site of transcription initiation, as alternative transcription start sites can dramatically change the magnitude of uORF repression.

## Results

### FACS-uORF determines the effects of thousands of AUG and non-AUG uORFs on protein expression

To evaluate the impacts of natural yeast uORFs on gene expression, we used FACS-uORF (Lin et al., 2019; Figure 1A). FACS-uORF simultaneously compares YFP expression from thousands of wildtype uORFs in endogenous transcript leaders to corresponding mutants in which the predicted uORF start codon has been mutated to a non-functional AAG. Our reporter library design included all transcript leaders less than 180 nucleotides long that contain at least one uAUG from *S. cerevisiae* (1,524 uORFs) and *S. paradoxus* (1,206 uORFs), as well as all *S. cerevisiae* non-AUG uORFs (540 uORFs) that we previously identified using ribosome profiling (Spealman et al., 2017). After removing low-frequency plasmid constructs with highly variable YFP levels (Methods, Figure S1), 1,689 unique AUG and 349 non-AUG uORFs remained with confident measurements of activity. To our knowledge, this represents the largest panel of natural uORFs assayed to date.

**Figure 1.**
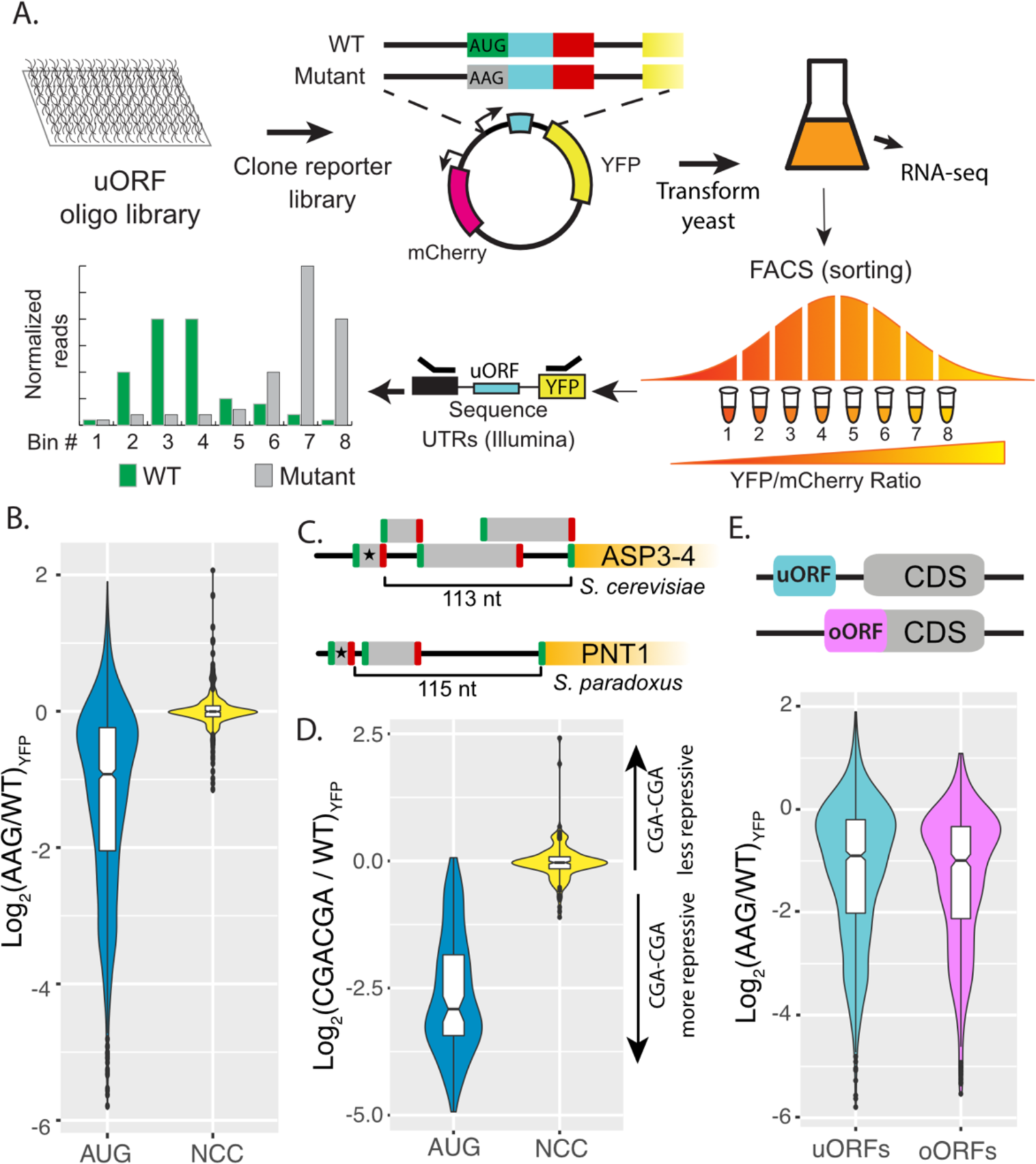
Regulatory impacts of 2,038 yeast uORFs. (A) A custom library of pairs of uORF-containing and uORF-mutant transcript leaders was cloned into reporter plasmids. Yeast were transformed with the reporter library and FACS sorted on the YFP / mCherry ratio. Plasmids extracted from the resulting FACS bins were sequenced to measure the expression levels of each reporter. (B) Comparison of AUG-and NCC-uORF activities. The log-fold change resulting from mutating uORF start codons is plotted. AUG uORFs are much more repressive than NCC-uORFs, though some AUG uORFs enhance expression. (C) Examples of complex transcript leaders with two enhancer uORFs (stars). (D) Insertion of CGACGA stalling dicodons increases repression from all AUG-uORFs and most NCC-uORFs, supporting their translation. (E) Overlapping AUG-uORFs (oORFs) are slightly more repressive than discrete uORFs.

We first evaluated the effects of AUG and non-AUG uORFs on reporter protein expression (Fig 1; Table S1). Most (1,222 of 1,689; 72%) AUG-initiated uORFs significantly altered YFP expression (Fig 1B). The vast majority (95%) of these functional uORFs were repressors, causing a 2.8-fold median decrease in expression. We next evaluated the 66 enhancer AUG-uORFs, as uORFs with enhancer activities are uncommon. Most (37) could be explained by gene annotation errors or alternative mechanisms (Figure S2). For example, the two strongest “enhancer” uORFs were preceded by a U, such that the (U)A<u>U</u>G -> A<u>A</u>G mutation used created UA<u>A</u> stop codon that converted unannotated N-terminal extensions into uORFs (Spar_9:277239-277245 and chrXV:423740-oORF). Several other enhancer uORFs had start codons preceding another AUG (<u>AUG</u>NAUG, e.g. uORF chrXII:73376-73397), such that the upstream AUG -> AAG mutation places -3A in the Kozak regions of the next AUG uORF, and are likely false positives resulting from activation of downstream uORFs (Table S2). After removing suspected false positives from further study, the remaining 29 enhancer uORFs increased expression from 1.15-fold to 4-fold, with a median increase of 1.7-fold. Most (26) were in multi-uORF transcript leaders, such that their translation might alter the use of other uORFs. Indeed 12 enhancer uORFs were upstream of other uORFs, reminiscent to the *GCN4* uORF1 enhancer that insulates against initiation at repressive downstream uORFs under stress (e.g. Fig 1C). Thus, while most uORFs reduce gene expression, a small number act as enhancers, potentially by insulating against the effects of other, more repressive uORFs.

In comparison to AUG uORFs, non-AUG uORFs were less likely to change expression (215/349, 62%, FET P = 8 x 10 ^-5) and had much smaller impacts, as mutating their start codons changed expression by 4-fold or less (median 10% change, WRT P = 2.2 x 10^-16; Fig 2A).

**Figure 2.**
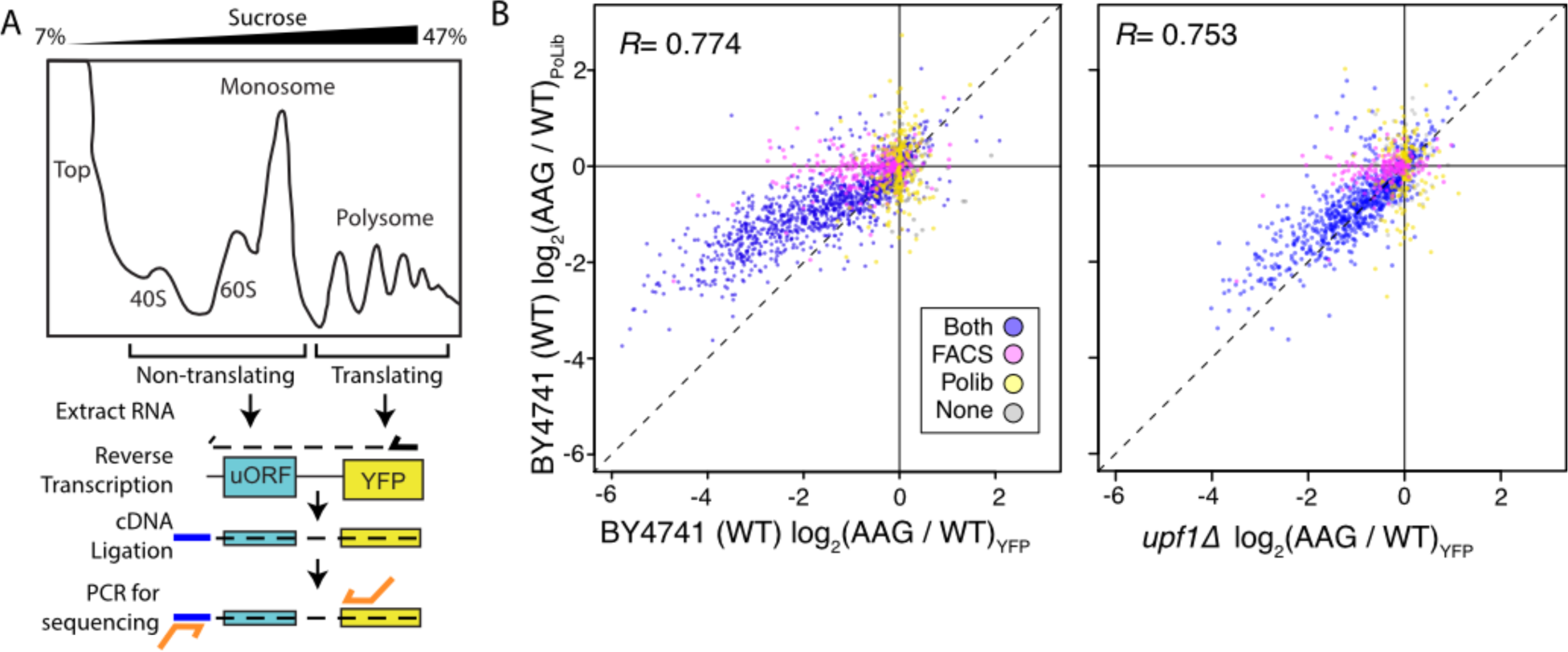
Massively parallel analysis of ribosome loading by Polysome Library Sequencing (Polib-seq) supports FACS-u ORF estimates of uORF function(A) The Poli b-seq assay system was developed to determine the impact of uORFs on ribosome loading. Polysome extracts were prepared from wildtype (BY4741) yeast strains expressing FACS-u ORF library 2 and separated on a 7 - 47% sucrose gradient by untracentrifugation. RNA was extracted from polysomal (2+ ribosome) and non-polysomal (40S, 60S, and monosome) fractions, and reporter constructs were quantified by targeted RNA-seq.The percent of polysomal reads was compared for each wildtype / AAG mutant uORF pair to determine the impact of each uORF on ribosome loading. (Bl Comparison of FACS-u ORF (x axis) and Poli b -seq (y axis) estimates of uORF function, both given as the log -trans fo rmed difference between wildtype and AAG mutant uORF. uORF impacts on ribosome loading are positively correlated with impacts on protein expression in wildtype yeast (left), and the correlation is stronger in the *upf1*Δ strain (right).

Non-AUG uORFs were eight times more likely to have enhancer activity than AUG-uORFs (102 / 215, 47%, FET P < 2.2x10^-16). A manual evaluation of thirty non-AUG uORFs that increased expression by at least 25% found six were located inside AUG uORFs, such that their mutation may alter the AUG uORF function. Similarly, four of sixteen non-AUG uORFs that decreased expression by at least 25% were nested inside AUG uORFs. The location of these non-AUG uORFs inside AUG uORFs complicates interpretation of their functions by start codon mutation. In summary, these results indicate that non-AUG uORFs have milder impacts on expression, consistent with inefficient initiation at non-AUG codons.

We previously showed that insertion of the CGACGA dicodon, which stalls elongating ribosomes (Letzring et al., 2013), increased repression from the yeast *YAP1* uORF (Lin et al., 2019). We reasoned that CGACGA insertion should similarly increase repression of other translated uORFs. To test this, we used FACS-uORF to compare YFP expression from 100 significant AUG- and 164 non-AUG uORFs with and without CGACGA insertions (Fig 1D; Table S3). The dicodon insertion made nearly all AUG-uORFs (99%), and most non-AUG uORFs (59%, BET P=0.0235), more repressive. These results support the active translation of AUG- and most non-AUG uORFs, both enhancers and repressors. However, the significant regulatory effect of mutating non-AUG uORF start codons to AAG may not always result from a loss of translation initiation, as a stalling dicodon did not cause repression from a considerable fraction of non-AUG uORFs.

We next considered the regulatory impact of uORFs that <u>o</u>verlap the main gene ORF (oORFs). Previous analyses of metazoan ribosome profiling data suggested that oORFs are more repressive than uORFs that terminate in transcript leaders (Chew et al., 2016; Johnstone et al., 2016). However, ribosome profiling may not accurately evaluate uORF regulatory effects because it captures noisy snapshots of ribosome occupancy over short sequence regions. By assaying the expression of wildtype and uORF-mutant reporter plasmids, we directly compared the regulatory effects of uORFs and oORFs in yeast. Considering all AUG-uORFs, we found that oORFs are ∼ 10% more repressive than uORFs (Fig 1E; WRT P = 0.02432), and this difference decreases when considering only AUG uORFs that have significant impacts on expression. Thus, our results show that oORFs are only slightly more repressive than uORFs in yeast.

### Consistent uORF impacts on protein levels and translation efficiency

By assaying steady-state protein levels, FACS-uORF measures aggregate changes in mRNA transcription, stability, and translation efficiency. As such, it is possible that the mutations used to inactivate uORF start codons might also affect reporter construct transcription and decay, rather than altering translation efficiency. This is a particular concern for apparent enhancer uORFs, which decrease YFP expression after start codon mutation. To validate the effects of uORFs on mRNA translation we developed a second MPRA for polysome loading, called <u>Po</u>lysome <u>Lib</u>rary <u>seq</u>uencing (PoLib-seq Fig 2A; Table S4). PoLib-seq involves sucrose gradient fractionation of polysome extracts from yeast carrying the reporter library, followed by directed RNA-sequencing to estimate the impact of uORFs on ribosome loading. PoLib-seq and FACS-uORF estimates were positively correlated, indicating general agreement between these two complimentary assays. Importantly, both enhancer uORFs and repressor uORFs identified from FACS-uORF had similar effects on ribosome loading as measured by PoLib-seq (Figure 2B). Of the 1,216 significant repressors of YFP found by FACS-uORF, 1,068 repressed ribosome loading in PoLib-seq measurements (88%, BET p < 2.2 * 10^-16). Similarly, of the 120 AUG and non-AUG FACS-uORF YFP enhancers, 86 increased ribosome loading in PoLib-seq (72% p < 2.3 * 10^-6). To further evaluate this, we performed FACS-uORF using a strain in which NMD is eliminated (*upf1Δ*). The slope of the linear regression between FACS-uORF and PoLib-seq results is closer to 1 in *upf1Δ* than in wildtype yeast (Fig 2B, p = 3.6 * 10^-31), indicating that PoLib-seq captures the translational effect of uORFs independent of NMD. Thus, the FACS-uORF and PoLib-seq results were generally consistent, underscoring the regulatory impact of these uORFs on mRNA translation. However, because PoLib-seq results were noisier than FACS-uORF results (Figure S3), we used FACS-uORF data for the remainder of this study.

### Dissecting the contribution of NMD in uORF repression

To examine the influence of NMD on uORF repression, we next compared uORF activity in wildtype and *upf1Δ* yeast (Fig 3A; Table S5). In general, repressor uORFs were *less* repressive in the *upf1Δ* strain, including 92% of AUG-uORFs and 59% of non-AUG uORFs. The decrease in repression that was observed among strong inhibitory non-AUG uORFs (Fig 3A inset) indicates that at least some non-AUG uORFs induce NMD. Since uORF-induced NMD directly impacts RNA abundance, we used targeted RNA-seq to compare the effects of uORFs on RNA levels. In wild-type yeast, 54% of the variance in uORF effects on YFP protein levels could be explained by differences in RNA abundance (Fig 3B). Notably, this correlation was essentially eliminated in *upf1Δ*, suggesting uORF-induced NMD plays a prominent role in reporter repression. (Fig 3B). By comparing the magnitude of uORF repression in wildtype (both translation and NMD) and *upf1Δ* (translation only), we quantified the contribution of NMD (%NMD; Fig 3C) for 431 significant repressor AUG-uORFs for which data were available from both strains. Across these repressive uORFs the median percentage of NMD is 35%. Thus, we estimate that roughly one third of yeast uORF repression is due to NMD.

**Figure 3.**
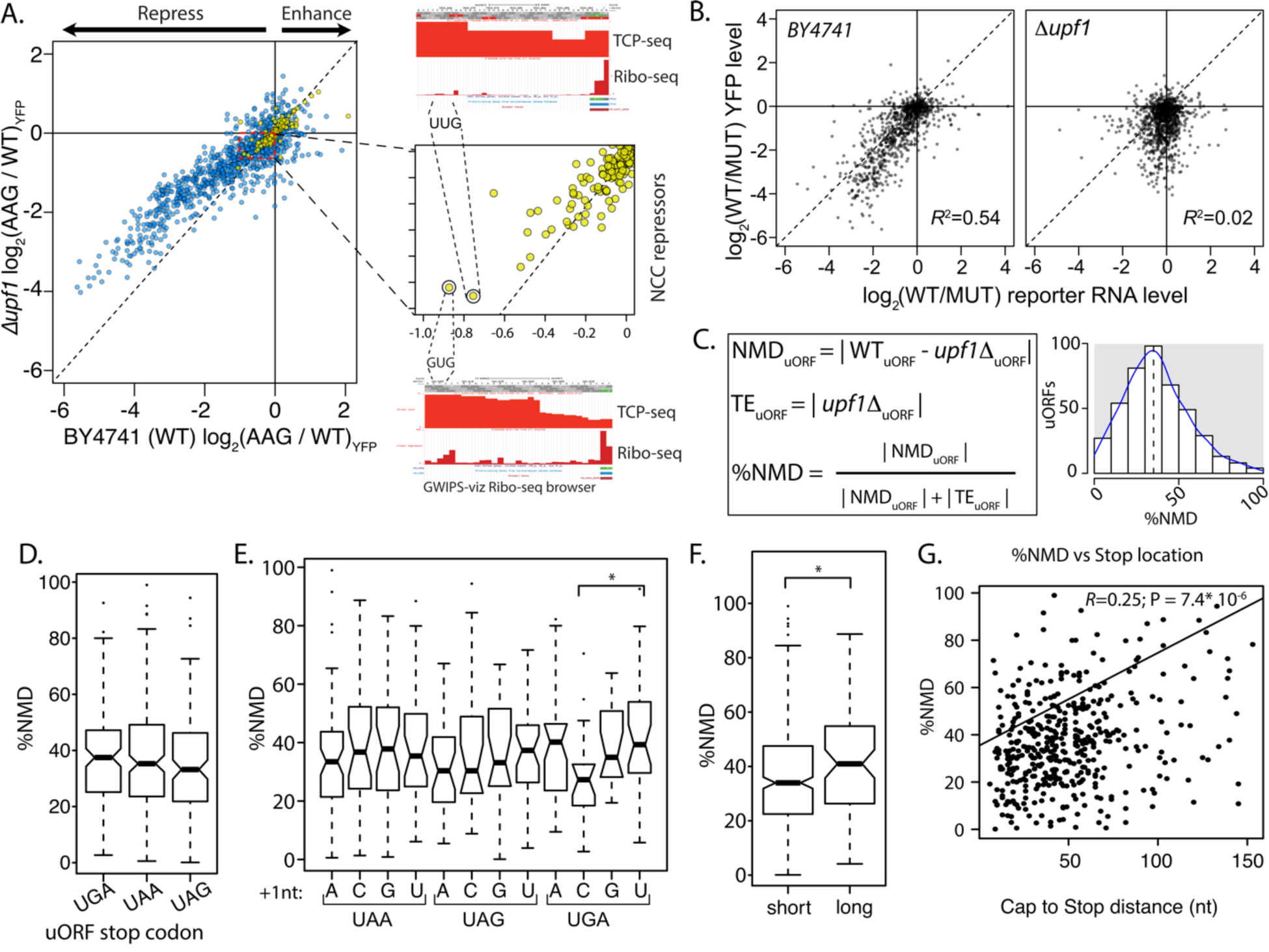
The role of nonsense mediated decay in uORF activity. (A) Scatter plot compares the regulatory impact of mutating uORF start codons in wildtype (x-axis)and *Δupf1* mutant yeast defective for nonsense mediated decay (left). Most AUG (blue), and some NCC-uORFs (yellow) were less repressive in a *Δupf1* strain, compared to wildtype . A zoomed view of NCC-uORF NMD effects is shown, highlighting two relatively strong NCC-repressor uORFs. Compos ite Ribo-seq and TCP-seq profiles from the GWIPS-viz browser supporting translation of these uORFs are shown (right). (B) Comparison of uORF impacts on mRNA and protein expression levels are shown for wildtype (left) and *Δupf1* (right) yeast strains. *Δupfl* decouples uORF impacts on transcription and translation . (C) Calculating the relative regulatory impact of uORFs on mRNA stability through NMD (NMD_u0RF_) and on protein translation efficiency (TE_u0RF_).The relative contributions of NMD to uORF activity (%NMD) is plotted for all assayed uORFs, showing a wide range of NMD induction . (D through G) Association of uORF features with %NMD.Termination at UGA stop codons induces more NMD than other stop codons (D), however UGA!: stop codons, which are known to allow read-through, have lower o/oNMD. Long uORFs induce more NMD than shorter uORFs, and uORFs that terminate far from transcript leader caps induce more NMD than those that terminate close to 5’ caps.

Our dataset provides a unique opportunity to examine the features associated with variation in uORF induction of NMD. NMD is induced through inefficient translation termination, and termination efficiency varies among the three stop codons (UAA∼UAG>>UGA) (Bonetti et al., 1995). Previous work suggested the extent of NMD induced by PTCs in coding regions depended on the stop codon identity (Keeling et al., 2004). Consistent with this, we found median %NMD was higher for uORFs terminating with UGA (37.5%) than those terminating with UAA (35.3%) or UAG (33.2%) (Fig 3D). The first nucleotide after the stop codon is known to influence termination through base-stacking interactions with rRNA (Brown et al., 2015). The %NMD varied among all 12 stop+1 sequences (Fig 3E). Although this was not generally significant (one way ANOVA), the large variation among UGAN stop contexts approaches significance, with UGAC having lower %NMD (29.7%) than UGAU (41.3%) (Fig 3E; Kruskal-Wallis test P = 0.061). UGAC stop codons have been shown to allow stop codon read-through at rates higher than any other stop codon (Cridge et al., 2018; Namy et al., 2001). In Summary, stop codon sequence context appears to influence the propensity for NMD by uORFs.

We reasoned that post-termination reinitiation at uORFs might also protect mRNA from NMD. As ribosomes more efficiently resume scanning after termination of short uORFs than long ones (Gunišová et al., 2017), we investigated the relationship between %NMD and uORF length. As expected, short uORFs (<=12 amino acids) exhibited lower %NMD than long uORFs (> 12 amino acids) suggesting that termination leading to reinitiation reduces the propensity for NMD (Fig 3E; Wilcoxon Rank-sum test P = 0.011). Unexpectedly, the location of the stop codon relative to the transcript leader cap was positively correlated with the %NMD, such that uORFs that terminate further from the cap were more likely to induce NMD than those that terminate adjacent to the cap (Fig 3F; R^2 = 0.065; P = 7.44x10^-8). Together, these results suggest that the propensity for uORFs to induce NMD depends on termination efficiency, uORF length, and uORF stop codon position in the mRNA transcript leader.

### Features that control uORF regulatory function

We next evaluated how uORF features affect their regulatory functions. The primary feature currently evaluated when examining uORF activity is the strength of the start codon Kozak context. In yeast, a strong Kozak context is typically rich in adenosine, particularly at the -3 position (Li et al., 2019). This characteristic “strong” Kozak context was only observed among the six most repressive uORFs (32-fold or more repressors). It was entirely absent for uORFs that decreased expression less than 16-fold and for enhancer uORFs (Fig 4A). This suggests that other features may influence the activity of most yeast uORFs. Intriguingly, we found that the relative position of uORF start and stop codons correlated with differences in regulatory activity. Start codons located near the transcription start site were more repressive than those positioned further downstream, while stop codons further from the main ORF (in this case YFP) start codon were less repressive (Fig 4B-C). These results suggest uORF location plays a more prominent role in determining regulatory function than has been previously appreciated.

**Figure 4.**
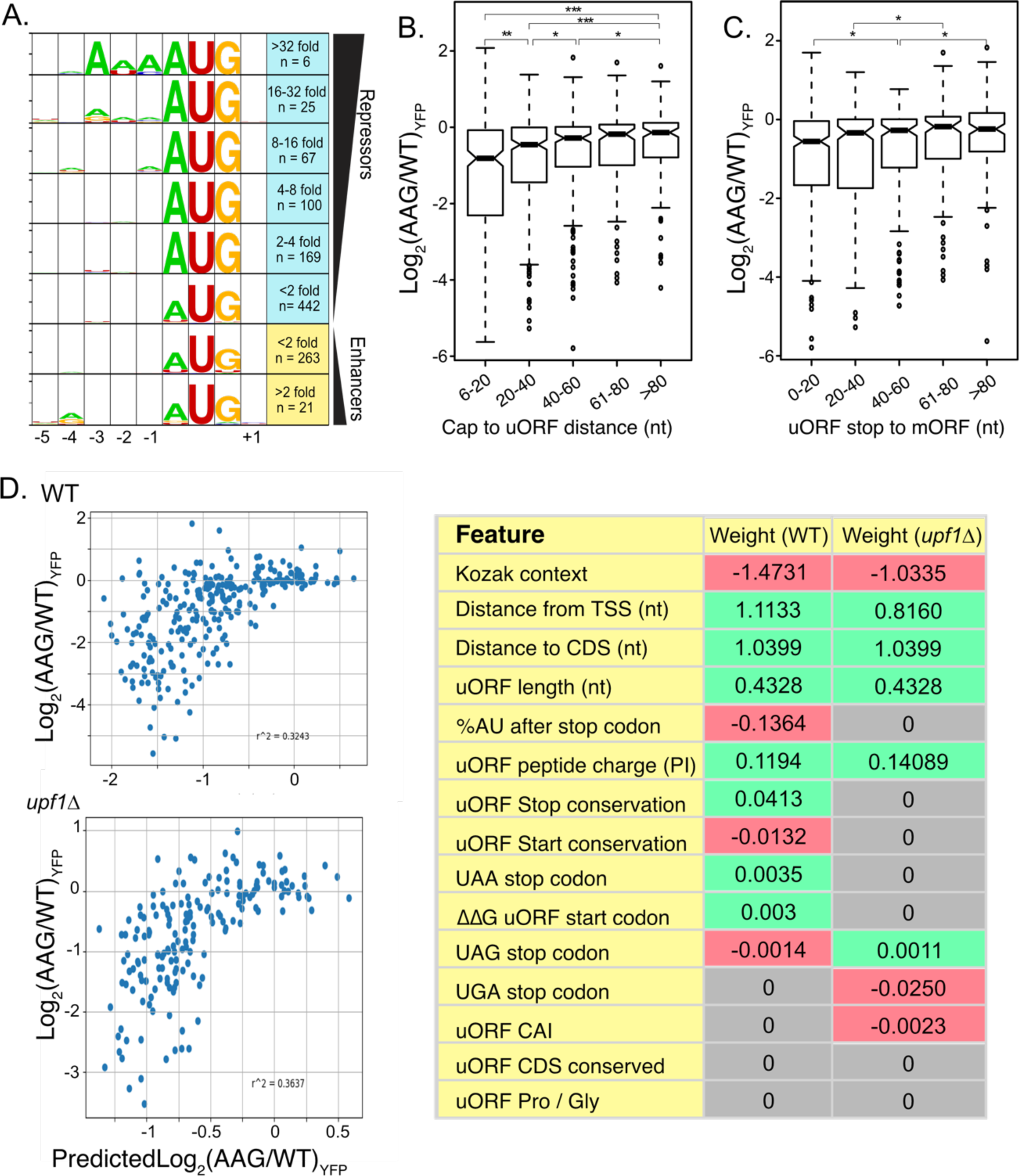
Elastic Net Regression modeling of uORF activities in wildtype and UPF1-delta strains. **A.** Weblogo consensus sequences in the Kozak region surrounding uORF start codons, separated by regulatory effect. **B.** Boxplot of regulatory effects for uORFs binned on distance between the 5’ cap and the uORF start codon. uORFs starting before 6 nucleotides were removed to avoid the A-rich region immediately after the reporter transcription start site. **C.** Boxplot of regulatory effects for uORFs binned by distance between the stop codon and the YFP start codon. **D.** Scatterplot comparing uORF activities predicted by the elastic net regression model (x-axis) with those observed from FACS-uORF (y-axis) in wildtype yeast. The model explains roughly 1/3 the variance in uORF activity. The table at right lists features selected by the ENR and their corresponding weights. Notably, uORF start and stop codon location features are together more impactful than Kozak sequence strength. Similar results for ENR modeling of uORF activity in a upf1-deletion strain. In the absence of NMD, the sequence downstream of the stop codon is no longer significant, while the uORF codon Adaptation Index (uORF CAI) becomes significant.

Because many features of uORF sequence and location can correlate with their impact on gene expression, their relative influence can be difficult to disentangle. For example, uORF Kozak contexts could vary with their location due to differences in G/C content near main ORF start codons. We next used a machine learning approach to select features that influence natural uORF activity independently and quantify their effects. Based on our results, and prior work, we examined fifteen uORF features, including Kozak context, folding energy around the start codon, uORF position in the transcript leader, codon usage and peptide charge, uORF length, stop codon identity and sequence context, and start and stop codon conservation. Given the size of our dataset, we chose elastic net regression (ENR) to select and weight features that contribute to a linear regression model of natural uORF activity (see methods). We built ENR models of uORF activity from both wildtype and *upf1Δ* strains to further investigate the role of uORF features in translational repression and NMD.

Many of the uORF features selected by the ENR models were similar for wildtype and *upf1Δ* strains. As expected, start codon context was the most influential feature selected by the ENR in both, having a strong negative correlation with uORF activity. Surprisingly, two features describing uORF location - distance from the cap to the start codon and from the stop codon to the main ORF - were selected independently and had nearly as strong an influence on regulation as Kozak contexts (Fig 5). Both features had strong positive correlations with uORF activity, as uORFs initiating closer to the 5’ cap, or terminating closer to the main ORF start codon, were more repressive.

**Figure 5.**
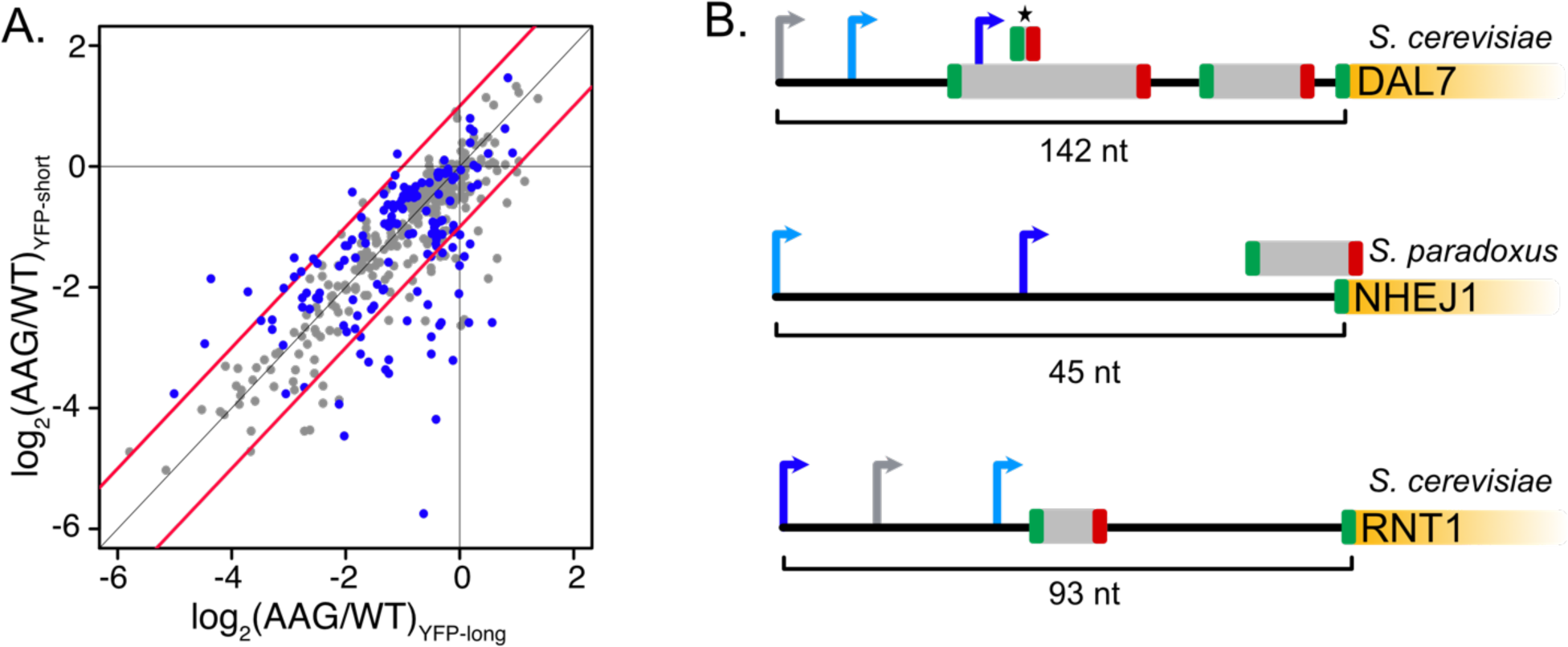
uORF activity depends on transcription start site usage. (A) Scatter plot comparing the activities of individual uORFs when located in longer (x axis) or shorter (y axis transcript leaders resulting from alternative transcription start sites. Significant changes in uORF activity are indicated as red points in the plot (t-test adjusted p < 0.05). Blue lines mark 2-fold difference in activity. In most cases, uORFs are more repressive in shorter transcript leaders, when located closer to the transcription start site and 5’ m7G cap. (B) Examples of uORFs with TSS location dependent activity. Arrows denote alternative transcription start sites. Blue arrows signify transcript leaders in which uORFs are more repressive, with dark blue indicating greater significance than light blue. Gray arrows indicate transcription start sites that were not significant. The asterisk shows the uORF whose repression varies with alternative transcription start site usage for DAL7.

uORF coding region features also contributed significantly, to the model, as shorter uORFs and uORFs encoding more negatively charged peptides were more repressive. However, these effects were relatively modest. Although ENR models from wildtype and *upf1*Δ strains were generally similar, AU-rich sequences downstream of uORF start codons were associated with increased uORF repression only in the wildtype ENR model. Together, the ENR models indicate that uORF Kozak context, position, length, and peptide charge are strong contributors to translational repression, while AU-rich downstream sequences may alter the propensity for NMD after uORF translation, consistent with historical reports of downstream sequence elements.

Finally, because the ENR models explain only a third of the variation in natural uORF activity, other features or non-linear relationships likely have additional influence in uORF function.

### Variation in uORF functions in alternative transcript leader isoforms

Yeast use alternative transcription start sites that create corresponding alternative transcript leader sequences in response to environmental stimuli (Arribere and Gilbert, 2013; Lu and Lin, 2019; Pelechano et al., 2013). We compared the activities (AAG / WT) of 333 uORFs in 470 alternative transcript leaders (Table S6). Strikingly, 116 uORFs differed significantly depending on the transcription start site (t-test, p.adjust <= 0.05; Figure 5A). In general, uORFs were more repressive when they were closer to the TSS. This result is consistent with our regression modeling, which identified uORF positions relative to 5’ cap and main ORF start codon as important features that impact uORF function. For example, the uORF from the *S. paradoxus NEJ1* gene is ∼60 fold repressive in the context of a transcript leader starting at 25 nucleotides upstream, and only 1.5-fold repressive when transcription is initiated a further 45 nucleotides upstream (Fig 6B). Both YFP mRNA and protein levels are extremely low in the former, suggesting that initiation at the downstream TSS is largely unproductive due to NMD and translational repression. A homologous uORF located in the *S. cereviaie NHEJ1* has similar dependence on transcription start site location, however its reporter expression values were too noisy for confident estimation. Similarly, a uORF in *S. cerevisiae DAL7* is not functional in the context of the longest (142 nt) transcript leader tested, but becomes a 1.3-fold repressor in an 124 nt leader and a 2.3-fold repressor when located just 11 nucleotides downstream of the 5’ cap.

**Figure 6.**
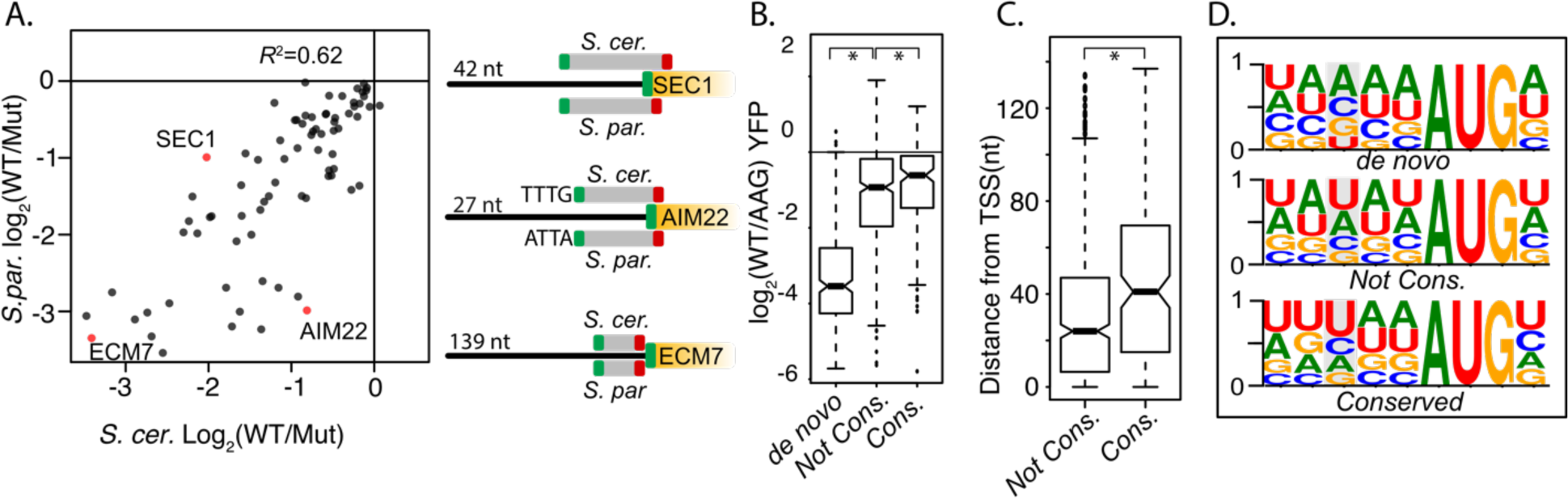
Evolutionary differences in uORF activities. (A) Scatter plot compares the activities of homologous uORFS from *S. cerevisiae* (x axis) and *S. paradoxus* (y axis) all assayed in S. *cerevisiae.* uORF activities are moderately correlated, suggesting relaxed selection on the magnitude of repression. (B) Boxplot depicts the magnitudes of uORF activities for de novo uORFs (mutation of NCC-uORFs to AUG-uORFs), conserved uORFs (Start codon PhastCons >= 0.5) and not-conserved uORFs (PhastCons <0.5). Natural uORFs are much less repressive than de novo uORFs, suggesting strong selective pressure to remove novel uORFs. Conserved uORFs are also significantly less repressive than non-conserved uORFs. (C) Conserved uORF start codons are farther from transcription start sites than non­conserved uORF starts. (D) Sequences in the Kozak consensus region of de novo, non-conserved, and conserved uORFs. The-3 position is highlighted in gray. Natural uORFs have a much lower frequency of-3 A than de novo uORFs, which likely contributes to their different magnitudes of repression.

However, there are exceptions to this general rule, as shown by a highly conserved uORF found upstream of *S. cerevisiae RNT1*. The *RNT1* uORF is a 1.3-fold repressor in its longest transcript leader context (93 nt), but shows weaker repression in two shorter transcript leader isoforms.

Together, these results show a striking dependency of uORF activity on the location of transcription initiation, underscoring the importance of position in uORF function.

### Evolutionary constraints on the magnitude of uORF activity

While previous studies have evaluated uORF conservation, these have generally not investigated conservation of uORF activity. Thus, we examined the activities of orthologous uORFs from *S. cerevisiae* and *S. paradoxus*, which last shared a common ancestor ∼ 5Mya (Scannell et al., 2011). Notably, our data compare these uORFs in their native homologous transcript leaders. After removing uORFs whose activity measurements were noisy (σ > 0.15) the direction of uORF regulation (repressor or enhancer) was almost entirely conserved (77/78; Table S7). However, conservation of the magnitude of regulation was less robust (*R*^2^ = 0.62, Fig 7A). For example, an *S. cerevisiae* oORF in *SEC1* was 1.5 to 2-times more repressive than its *S*. *paradoxus* homolog, potentially owing to a deletion in *S. paradoxus* that results in an earlier stop codon that shortens the oORF. In another case, an *S. paradoxus* oORF in the *AIM22* leader was approximately four-fold more repressive than its *S. cerevisiae* homolog, possibly due to the presence of more adenosines in its Kozak sequence. In contrast, the sequence and location features of the *ECM7* uORF were highly conserved, and both species uORFs had strong 10-fold repression. Thus, the magnitude of repression at many yeast uORFs has diverged substantially, though their functions as repressors are generally conserved.

We next divided uORFs by the average PhastCons Score (PCS) over their start codons into “conserved” (PCS > 0.5) and “non-conserved” (PCS < 0.5) groups. Consistent with predictions from vertebrate Ribo-seq studies (Chew et al., Johnstone et al) conserved uORFs were less repressive than non-conserved (Fig 7B). We also found that mutating NCC-uORF start codons to AUG created much more repressive “de novo” uORFs (Fig 7B; Table S8). These results suggest that newly emergent yeast uORFs tend to be strong repressors that are likely removed by natural selection. To investigate potential explanations for the decreased repression exhibited by conserved uORFs, we compared the features of “conserved” and ‘non-conserved” uORFs selected by our ENR modeling. Interestingly, conserved uORF start codons were located further from transcription start sites (Fig 6C) and had a lower frequency of adenosine at the -3 position of their Kozak contexts (Fig 6D) than their non-conserved counterparts. These results suggest that both sequence and position features are subject to selection, such that conserved uORFs tend to have features associated with milder impacts on expression.

## Discussion

Thousands of AUG and non-AUG uORFs have been identified from ribosome profiling studies in all major model organisms (Kearse and Wilusz, 2017). Despite concerns that cycloheximide and other drugs used in ribosome profiling may exaggerate occupancy on transcript leaders (Gerashchenko and Gladyshev, 2014; Kearse et al., 2019; Lareau et al., 2014), very few natural uORFs have been experimentally tested because traditional uORF assays are performed on a gene-by-gene basis (Wethmar et al., 2014). Here, we used two Massively Parallel Reporter Assay systems, FACS-uORF and PoLib-seq, to quantify the impact of thousands of yeast AUG- and non-AUG uORFs on reporter expression and translation efficiency in wildtype and NMD-deficient yeast strains. We leveraged the resulting data to investigate the importance of multiple uORF features on their regulatory functions. Our analyses shed new light on uORF functions and have multiple implications.

It has long been recognized that eukaryotic translation can initiate at non-AUG (so-called) near-cognate codons (Kearse and Wilusz, 2017; Kozak, 1991; Tang et al., 2004). Ribosome profiling studies also support the translation of many non-AUG uORFs in yeast (Brar et al., 2011; Eisenberg et al., 2020; Ingolia et al., 2009; Spealman et al., 2018). However, the few non-AUG uORFs that have been experimentally tested had relatively modest regulatory effects in reporter systems (Spealman et al., 2018; F Zhang and Hinnebusch, 2011). Our results show non-AUG uORFs have relatively mild impacts on gene expression. Furthermore, a considerable number of non-AUG uORFs whose mutation significantly changed reporter expression may not be translated, as insertion of the stalling CGA dicodon sequence did not reduce expression.

Thus, at least during log-phase growth, many non-AUG uORFs appear to have little influence on gene expression, consistent with the relatively low rate of translation initiation at near-cognate start codons (Kolitz et al., 2009; Takacs et al., 2011). As non-AUG uORFs have been predicted in many other species, our results suggest that most of them may also play limited roles in regulating expression. However, non-AUG uORFs with start codons in strong Kozak contexts with accompanying downstream mRNA structure may more have more substantial effects.

Although most uORFs are expected to repress expression, examples of enhancer uORFs have been identified in multiple species. Most notably, the short upstream uORFs in *GCN4* (yeast) and *ATF4* (metazoans) have been shown to cause stress-dependent upregulation of main ORF translation (Hinnebusch, 2005; Vattem and Wek, 2004). This upregulation results from delayed reinitiation under stress, such that PICs that resume scanning bypass more repressive uORFs. In our system, only two percent of uORFs increased expression. Notably, the oligonucleotide synthesis technology used to generate our reporter library is limited in length, such that the longest leaders that we tested were 180 nucleotides. While this included most yeast transcript leaders, longer leaders may have more enhancers. As such, our numbers may underestimate the frequency of enhancers. Because most of the enhancers we found were in multi-uORF transcript leaders, we propose that, much like *GCN4* uORF1, they function by reducing usage of other, more repressive uORFs. Although we performed our experiments under unstressed conditions, reinitiation might still allow leaky scanning past more repressive uORFs near the enhancer stop codon. In other cases, translation of enhancer uORFs might alter transcript leader structure or modulate scanning by upstream PICs. Regardless of the mechanisms used, our data indicate that uORFs rarely act as enhancers, at least under the conditions we assayed.

It is well known that uORF activity depends on the strength of start codon recognition. However, the influence of other factors has been less clear. Our computational modeling of a massive set of yeast uORFs identified the relative influence of these and other features on uORF activity. While the influence of Kozak context was strong, surprisingly, uORF location had an effect comparable in magnitude. Because PIC assembly involves unwinding the mRNA around the 5’cap, the increased repression we observed for uORFs that start near the cap might result from increased start codon accessibility. Cap-proximal uORFs might also sterically block the loading of additional PICs. We also observed stronger repression for longer uORFs and uORFs that terminate closer to the main ORF start codon. Such uORFs may permit less reinitiation at the main ORF, as ribosomes lose initiation factors after translating PICs that resume scanning would have less time to reacquire ternary complex before encountering the main ORF start codon. If so, our results suggest translation reinitiation is more common in yeast than currently appreciated (Gunišová et al., 2017). Future work is needed to evaluate these and other potential mechanisms underlying the importance of uORF location.

Despite the insights gained by our ENR modeling, a large amount of the variance among uORFs cannot currently be explained. This suggests that other features may have important influences on uORF activity. For example, previous work found that structural accessibility of uORF start codons, as measured by SHAPE probing, led to accurate estimates of uORF activity for the human antitrypsin-alpha gene (Corley et al.). Based on this work, it was proposed that ribosomes often shunt past uORFs whose start codons are occluded in stable RNA structures (Mustoe et al., 2018). Although intriguing, this model seems to negate the role of helicases (e.g. DED1 and eIF4A) which unwind such structures during PIC scanning (Guenther et al., 2018; Sen et al., 2019, 2015; Sharma and Jankowsky, 2014). While we included predicted energies of unwinding around the start codon, our purely computational structural predictions contributed only minimally to our models. Given the large amount of uncertainty surrounding computational estimates of RNA structure, we expect that detailed measurements of structural accessibility would be beneficial to future modeling.

By comparing the magnitude of uORF repression in wildtype and *upf1*Δ yeast, we found the contribution of NMD to uORF repression (%NMD) ranges from 0 to 100%, with a median of 35%. Our work also provides insight into how uORF features impact their ability to induce NMD. Reinitiation after uORF translation can protect human mRNA from NMD (Lindeboom et al., 2016; Zhang and Maquat, 1997), and short uORFs are known to reinitiate more efficiently than long ones (Gunišová et al., 2017, 2016) likely due to increased retention of initiation factors (Bohlen et al., 2020; Wagner et al., 2020). Consistent with this, we found short uORFs induce less NMD than long uORFs. We also found NMD induction was lowest for uORFs terminating in UGA<u>C</u>. Since this stop codon context allows high rates of readthrough, it is possible such readthrough protects mRNA from uORF-induced NMD consistent with previous work (Keeling et al., 2004). Notably, AU-rich elements downstream of stop codons were associated with stronger uORF repression in wildtype, but not *upf1*Δ yeast. Such sequences may function as NMD-inducing *cis*-elements formerly thought to promote NMD by binding Hrp1p (Culbertson and Leeds, 2003; Czaplinski et al., 1999; González et al., 2000; Peltz et al., 1993; Ruiz-Echevarría et al., 1998). Thus, our work supports significant roles for many sequence features in determining uORF NMD induction.

We also investigated the relationship between uORF location in transcript leaders and NMD induction. Previous work using synthetic NMD targets found that NMD was more efficient when stop codons were located closer to the 5’ end of the *PGK1* coding sequence (Cao and Parker, 2003). We found the opposite relationship among uORF stop codons, such that stop codons were less likely to induce decay the closer they were located to the 5’ cap. Although these results at first appear discordant, it is possible circularization of the mRNA in the “closed-loop” model positions cap-proximal start codons closer to the poly-adenosine tail which could increase the efficiency of uORF termination. Thus, our work implicates uORF position is also an important feature impacting the efficiency of NMD.

Eukaryotic transcription often initiates at alternative sites, even in the relatively simple yeast (Lu and Lin, 2019; Pelechano et al., 2013; Zhang and Dietrich, 2005b). Such alternative transcription start sites alter the translation efficiency of their downstream genes (Arribere and Gilbert, 2013; Rojas-Duran and Gilbert, 2012; Zydowicz-Machtel et al., 2018). However, the extent to which uORF function depends on transcription start site usage has not been previously evaluated to our knowledge. Our results indicate that uORF activity often varies with alternative transcription start site usage. The dependence of uORF activity on transcript start sites may impart different translation efficiency and turnover rates for alternative transcript isoforms. This was broadly consistent with our modeling data, in that uORFs were more repressive when present in transcript leaders that initiated closer to the uORF start codon. If uORF start codons near the 5’ cap are more structurally accessible, uORFs that initiate further downstream may be bypassed by ribosomal shunting, as occurs in the Cauliflower Mosaic Virus uORF and has been proposed recently to be more common (Corley et al., 2017; Mustoe et al., 2018; Ryabova and Hohn, 2000). Alternatively, PIC scanning may become more processive over time, such that more distal start codons are more easily skipped. Regardless of the underlying mechanisms, the dependence of uORF activity on location implies that caution must be taken when studying uORFs using ribosome profiling data because current methods do not connect uORF occupancy with specific transcript isoforms.

Finally, our study included the first direct comparisons of gene regulation from homologous uORFs from two species in their corresponding, native transcript leaders. We found the magnitude of uORF regulation varied substantially for many uORFs, with some examples consistent with changes in uORF features. Our results also show a conserved uORF in yeast *ECM7* confers strong repression in both *S. cerevisiae* and *S. paradoxus*. Although previous work reported the uORF had no impact on regulation (Zhang and Dietrich, 2005a), this may have been due to non-physiological transcription initiation site usage. The luciferase reporter used in the previous study was driven by a different promoter which may have initiated transcription at alternative site(s). Earlier studies comparing ribosome profiling data from zebrafish, mouse, and human cell lines indicated that conserved uORFs tended to have weaker Kozak sequences than non-conserved uORFs (Chew et al., 2016; Johnstone et al., 2016). They proposed that this allowed conserved uORFs to be activated by *trans*-acting factors. In addition, conserved uORFs were associated with smaller decreases in ribosome occupancy on downstream main ORFs. By directly assaying uORF effects, we found that conserved uORFs indeed have more modest effects on gene expression than non-conserved uORFs. We also found uORFs with conserved start codons had weaker Kozak sequences and were located further from the transcription start site. These results suggest that both uORF Kozak context and location are subject to purifying selection, underscoring the importance of uORF position we identified by computational modeling.

In providing the first, to our knowledge, high-throughput functional analysis of natural uORF regulatory activities, our work defines the range of uORF activity and reveals location is as important as Kozak strength in determining yeast uORF functions. Recent work has also highlighted the large number of human uORFs (Barbosa et al., 2013; Calvo et al., 2009; Lee et al., 2020; McGillivray et al., 2018; Wethmar et al., 2010). While many aspects of translation initiation are deeply conserved, the mechanisms involved in transcript leader scanning may differ substantially in human cells and tissues. Thus, studies using similar methods in human cells are needed to characterize human uORF regulatory functions, and further understand their potential involvement in cellular stress responses.

## Data Accessibility

High-throughput sequencing data have been submitted to NCBI under Bioproject accession number PRJNA639207

## Acknowledgements

This work was supported by the National Institute of General Medical Sciences to C.J.M. (grant R01GM121895) and to J.W. (grant R01GM028301).

## Methods

### FACS-uORF library construction

The reporter plasmid was modified from *pGM-YFP-mcherry* (Lin et al., 2019)by replacing the GPMI promotor with an *ENO2* promotor. The *ENO2* promotor transcription start site is highly stringent (Spealman et al., 2018), allowing for highly consistent transcription starts sites in the transcript leader library. The *ENO2* promotor was inserted upstream of YFP by amplifying the ENO2 sequence from *S. cerevisiae* genomic DNA, using the primers Eno2_SalI-F and Eno2_AvrII_R1. The reverse primer introduced an AvrII restriction site in *ENO2* for use during subsequent cloning steps. The PCR product was then amplified using the primers Eno2_SalI-F and Eno2_XmaI_R2 to add additional Eno2 sequence and an XmaI site. The ENO2 PCR product was digested with SalI and XmaI, and ligated into the vector, *pGM-YFP-mCherry* resulting in the plasmid construct *pGM-ENO2-YFP-mCherry*.

The library was synthesized as two pools of 130- and 210-mer oligonucleotides (Agilent Technologies). Each oligonucleotide in the pool was designed to have a common *ENO2* promotor sequence and AvrII site upstream, and a yellow fluorescent protein (YFP) sequence downstream of the transcript leader so that the pool of oligos could be amplified and cloned into a dual fluorescent reporter. The oligo pool was provided as a 10 pmol pellet and was dissolved in 100 μl of TE. The primers Eno2_lib_F1and FACS-uORF-YFP-R, were used to amplify 30 μl (3 pmol) of the oligo pool in a 400 μl reaction (split up in 50 μl aliquots) containing 1X Herculase II reaction buffer, 0.4 mM each dNTP, 0.25 μM each primer, and 16 μl Herculase II Fusion DNA Polymerase (Agilent Technologies). The reaction conditions for the PCR reaction were 95 °C for 1 minute, followed by 10 cycles of 95 °C for 20 seconds, 55 °C for 20 seconds and 68 °C for 20 seconds, and then one cycle of 68 °C for 4 minutes. The PCR product was purified using AMPure XP beads (Agencourt) and resuspended in 20 μl of water. Half of the PCR product (9 μl) was used as a template for a second round of amplification, using the primers Eno2_lib_AvrII_F2 and FACS-uORF-YFP-R in a 400 μl reaction containing 1X Q5 polymerase reaction mix, 0.2 mM each dNTP, 0.5 μM each primer, and 8 units of Q5® High-Fidelity DNA polymerase. The amplification conditions were 98 °C for 30 seconds, followed by 10 cycles of 98 °C for 10 seconds, 55 °C for 20 seconds and 72 °C for 30 seconds, and then one cycle of 72°C for 2 minutes. The PCR product was purified over a Zymo Clean and Concentrator column (Zymo Research), digested with AvrII and BglII, and ligated into *pGM-ENO2-YFP-mcherry.* The ligated plasmids were then transformed into competent *E. coli*. Positive transformants were selected on LB plates containing ampicillin. In order to maintain the complexity of the library, a total of 100,000 positive colonies were scraped off 45 10 cm plates and pelleted. The plasmid library was extracted using a QIAGEN plasmid maxiprep column (Qiagen) and resuspended in 1 ml of TE.

### Section 1.01 Yeast transformation

Competent cells for each yeast strain were prepared using the Frozen-EZ Yeast Transformation II Kit™ (Zymo Research) according to the manufacturers’ instructions. For each strain, 400 μl of competent cells were mixed with 2 μg of each plasmid library and incubated shaking at 30 °C for 2 hours. To test the transformation efficiency, 10 μl of cells were plated on minimal media plates lacking uracil and incubated for 48 hours at 30 °C. The remaining cells were added to 30 ml of URA-media and incubated overnight shaking at 30 °C. The next day, the cells were added to 200 ml of URA-media and incubated overnight shaking at 30 °C. To ensure that at least 100,000 individual cells were transformed, the number of colonies from 10 μl of transformed cells was counted. For all strains, the total number of total clonal transformants ranged from 500,000 to 1 million. Glycerol stocks were made from each uORF library in each strain, by pelting cells from 10 ml of overnight cultures.

### Fluorescence activated cell sorting (FACS)

For each strain of yeast, one glycerol stock tube (10 ml of overnight culture) of each uORF library was culture was added to 200 ml of URA-media and grown shaking overnight at 30 °C. The cells were then restarted in 50 ml of URA-media at an OD600 of 0.1 - 0.2 and grown shaking 30 °C to an OD600 of 0.7, prior to FACS. Just prior to cell sorting, 12 ml of cells were pelleted and flash frozen for later RNA extraction, and 1 ml of cells were pelted and frozen for DNA extraction. The cells were sorted into nine bins, based on the ratio of YFP to mCherry, on a FACSVantage Digital Cell Sorter. For each bin, 100,000 cells were deposited into culture tubes containing 5 ml of URA-glucose media, and grown overnight shaking at 30 °C. To verify that the cells were sorted into the correct bins, and to later adjust the bin values to the ratio of YFP and mCherry, the ratio of YFP over mCherry for each bin was measured on a Tecan M1000 plate reader.

### Sequencing Library preparation

For the RNA sequencing libraries, 5 μg of total RNA was subjected to DNase treatment, re-extracted with acid phenol, ethanol precipitated over a column, and resuspended in nuclease free water. 2 μg of DNase treated total RNA was reverse transcribed with a RT primer (FacsuORF-RT) that anneals to the YFP sequence downstream of subsequent PCR primers. The cDNA was ethanol precipitated and resuspended in 10 μl of nuclease free water. The adapter, “Fuorf_RNA_adapter” was ligated to the 3’ end of 10 μl of cDNA in a 20 μl reaction at 65 °C for 1 hour using Thermostable 5’ App DNA/RNA ligase (New England Biolabs). The reaction was inactivated at 95 °C for 3 minutes and 2 μl of the reaction was used as a template for PCR for sequencing library generation as described below.

The plasmid DNA libraries from the unsorted library and from each bin were prepared by three rounds of PCR. Using primers that anneal to the ENO2 promoter (Eno2_lib_F1), and YFP sequences (FACs-uORF-YFP-R) 8 cycles of PCR were performed in a 20 μl reaction using 10 ng of plasmid library (or 2 μl of cDNA, as described above) as a template. The first PCR reaction was purified using 1.5X AMPure XP beads (Agencourt) and resuspended in 10 μl of water and added to a second reaction where 6 cycles of PCR was performed using primers that included varying numbers of random bases to stagger the sequencing reads (FuORF_2_DNA_N0-N7_F, and FuORF_2_DNA_N0,2,6_R). The second PCR was purified using 1.5X AMPure XP beads (Agencourt) and resuspended in 50 μl of nuclease free water. Using 2 μl of the second PCR as a template, a third round of 15 PCR cycles was used to incorporate Illumina sequences and 6 nucleotide barcodes, using the primers RPF_F and RPF_R. All PCR amplifications were performed using a high-fidelity polymerase (Q5 DNA polymerase, New England Biolabs) to avoid PCR introduced sequence error.

### PoLib-seq

One glycerol stock tube (10 ml of overnight culture) of each uORF library was added to 200 ml of URA-media and grown shaking overnight at 30 °C. The cells were then restarted in 50 ml of URA-media at an OD600 of 0.2 and grown shaking 30 °C to an OD600 of 0.7 and harvested by vacuum filtration. The cells were scraped off of the filter and flash frozen in liquid nitrogen. The yeast were cryoground using a mortar and pestle in 1 ml frozen polysome lysis buffer (PLB, 10 mM Tris-HCl pH 7.5, 0.1 M NaCl, 30 mM MgCl2, 50 mg/ml heparin, 50 mg/ml cycloheximide). The ground frozen yeast were added to 1.5 ml fresh PLB in a 2 ml microcentrifuge tube and thawed on ice. Approximately 0.5 ml of 0.5 mm zirconia/silica disruption beads were added to the lysates and the lysates were ground by vortexing for 30 seconds and then cooled for 30 seconds on ice for a total of four grinding cycles. The lysates were cleared by centrifuging for 10 minutes at 20,000G at 4 °C. The OD254/ml were determined and the lysates were flash frozen in liquid nitrogen in 50 OD260 aliquots. Forty OD260 units of lysates were layered on 7–47% (w/v) sucrose gradients, centrifuged (4 hours, 4°C at 27, 000 rpm) using Beckman L7 ultracentrifuge. A Teledyne ISCO Foxy R1 density gradient fractionator was used to fractionate and analyze gradients with continuous monitoring at OD254. Ribosomal subunits, ribosomes, and polyribosomes were fractionated according to the manufacturer’s protocol, and appropriate fractions (“top fraction”, 40S, 60S, monosome, 2, 3, 4, and greater than 5 ribosomes) were collected. RNA from each fraction (“top fraction”, 40S, 60S, monosome, 2, 3, 4, and greater than 5 ribosomes) was extracted by two rounds of acid-phenol extraction, purified over RNA clean up and concentrator – 5 (Zymo Research) columns, and eluted in nuclease free water. RNA sequencing libraries were prepared from each fraction (see Sequencing library preparation).

### FACS-uORF and PoLib-seq sequencing and data analysis

FACS-uORF and PoLib-seq libraries were subjected to paired-end (2 x 150 cycle) Illumina sequencing (Novogene). Read pair data were merged and error corrected using FLASH (Magoč and Salzberg, 2011) using parameters “-z -O -t 1 -M 150”. ENO2 promoter plasmid sequence present in DNA libraries (FACS only) was removed from the resulting merged reads using cutadapt (Martin, 2011) with parameters “--trimmed-only -e 0.04 -g AGTTTCTTTCATAACACCAAGC “. The resulting trimmed reads from each FACS bin were separately counted for perfect matches to the designed library UTR sequences using a custom perl script (DNA-seqcount.pl). YFP reporter RNA library data were merged and error corrected using FLASH as above. The resulting merged RNA-seq reads were further processed with cutadapt parameters “--trimmed-only -e 0.04 -g AGTTTCTTTCATAACACCAAGCNNNN” to remove the 3’ Fuorf_RNA_adapter sequence ligated to cDNA during RNA library preparations. After this adapter trimming step, many resulting RNA-seq reads contained two random bases and a guanosine, “NNG”, preceding the designed transcription start site sequence “AAGC” from the ENO2 transcript leader. on their 5’ ends. These “extra” sequences likely reflect reverse transcription of the 5’ 7mG cap (the universal “G”) followed by the untemplated addition of two random bases during cDNA synthesis. Cutadapt was used to remove these sequences, with parameters “--trimmed-only -g ^NNGAAGC“ to identify 5’-capped reads and “--discard -g ^NNGAAGC” to identify uncapped reads. The resulting reads from each sample were counted for perfect matches to the designed UTR library using the custom perl script (RNA-seqcount.pl).

### FACS-uORF analysis

For each of the three FACS-uORF replicates, count data from the RNA-seq library, total plasmid library (“bin0”), and libraries from each of the eight FACS bins (bin1 - bin9) were normalized and processed as previously described (Lin et al., 2019) using a custom perl script (“FuORF-Tables-Maker.pl”). Briefly, the relative representation of DNA and RNA reads was calculated as the reads-per-million for each replicate, and relative transcript levels were estimated as RNA-rpm / DNA-rpm. YFP/mCherry expression levels for each bin were taken from TECAN readings of overnight sorted cultures (see FACS methods) and normalized to the highest value (bin8 = 1). Read count data from each bin were downsampled such that the proportion of total reads from each bin reflected the proportion of cells sorted into that bin. After these normalization steps, the average YFP value for each UTR in the library was calculated as follows: YFP / mCherry = (SUM(YFPbin * reads / bin) from bin 1 - 9)/total reads). The resulting output files contained estimates of RNA and YFP levels for each UTR. The three replicates were compared, and UTRs with noisy YFP measurements (standard deviation > 0.05, minimum of 50 normalized reads per UTR) were removed from further consideration (Figure S1). For each uORF, mean YFP values from wildtype and mutant UTRs were compared using the wilcoxon rank-sum test in each replicate, and the benjamini - hochberg correction was applied to control the FDR at 5%. uORFs with significant (FDR < 0.05) and consistent (same direction) effects on YFP levels were considered significant regulators. Statistical analyses were done using R (version 3.6.1).

### PoLib-seq analysis

PoLib-seq libraries were pooled to provide sequence read counts in proportion to the total RNA present in each sucrose gradient fraction (40S, 60S, monosome, 2, 3, 4, 5+) (Figure S4). To compensate for variations in manual fractionation of the replicate sucrose gradients, we pooled “translating” (2, 3, 4, and 5+ ribosomes) and “non-translating” (40S, 60S, and monosome) fractions separately for each replicate, similar to RNA-seq based polysome profiling analyses (Anota2seq, e.g. (Oertlin et al., 2017)). The “top” fraction was not used for analysis. After manual inspection, the monosome fraction of replicate 1 was downsampled by a factor of 0.8626 to ensure similar proportions of “translating” (45%) and “non-translating” (55%) reads from each of the two replicates (Figure S4). A 5,000 total read cutoff (capped and uncapped, replicates 1 and 2 summed across all fractions) was used for each wildtype / mutant uORF comparison. The impact of each uORF on ribosome loading was calculated as the fold change in the translated / untranslated ratio from the mutant UTR compared to the wildtype UTR using a custom perl script (PoLib-Tables-Maker.pl). uORFs with consistent (same direction) influences on expression were tested for significance using the cochran-mantel haenszel test, and an FDR of 5% was maintained after benjamini-hochberg correction of raw p-values. Statistical analyses were done using R (version 3.6.1).

### uORF Modeling and Feature Selection by Elastic Net Regression

Elastic net regression attempts to solve the feature-selection and shrinkage problems simultaneously with the introduction of both L1 and L2 regularization. The L1 parameter functions similarly to LASSO regression and selects features, and the L2 parameter minimizes the coefficients of predictors (Hastie et al., 2009). The model minimizes the objective function:

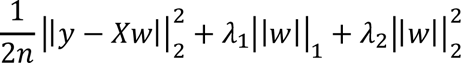

With and representing the L1 and L2 regularization parameters respectively. Regression was performed using the sklearn ElasticNet package, which takes two hyperparameters. The score penalty α corresponds to the magnitude of the regularization penalties, and the *l1_ratio* parameter () is used to control the relative values of the L1 and L2 penalties, such that:

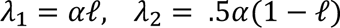

Fifteen features were selected from the data and were normalized by minmax. Test and training data were separated according to an 80/20 split. The hyperparameters α and were tuned manually to balance model performance and precision over 100 trials with randomly initialized training and test data. Model performance in this case refers to the R2 score of the model on the test set, and precision refers to the similarity of selected feature subsets between trials.

It has been previously shown that the strength of the Kozak sequence is strongly implicated in the translational regulation of uORFs. For this reason, single-feature linear regression on Kozak strength alone was used as a control to demonstrate the robustness of the model. For each of the 100 trials, the R2 score of the Elastic Net model against single-feature LR was determined, and p-values were computed using the Wilcoxon rank-sum test.

## Acknowledgements

This work was supported by the National Institute of General Medical Sciences (grant R01GM121895 to C.J.M. and R01GM028301 to J.W.).

## Data Availability

Raw sequencing reads have been deposited at NCBI SRA under project accession number PRJNA721222.

---

**Figure S1.**
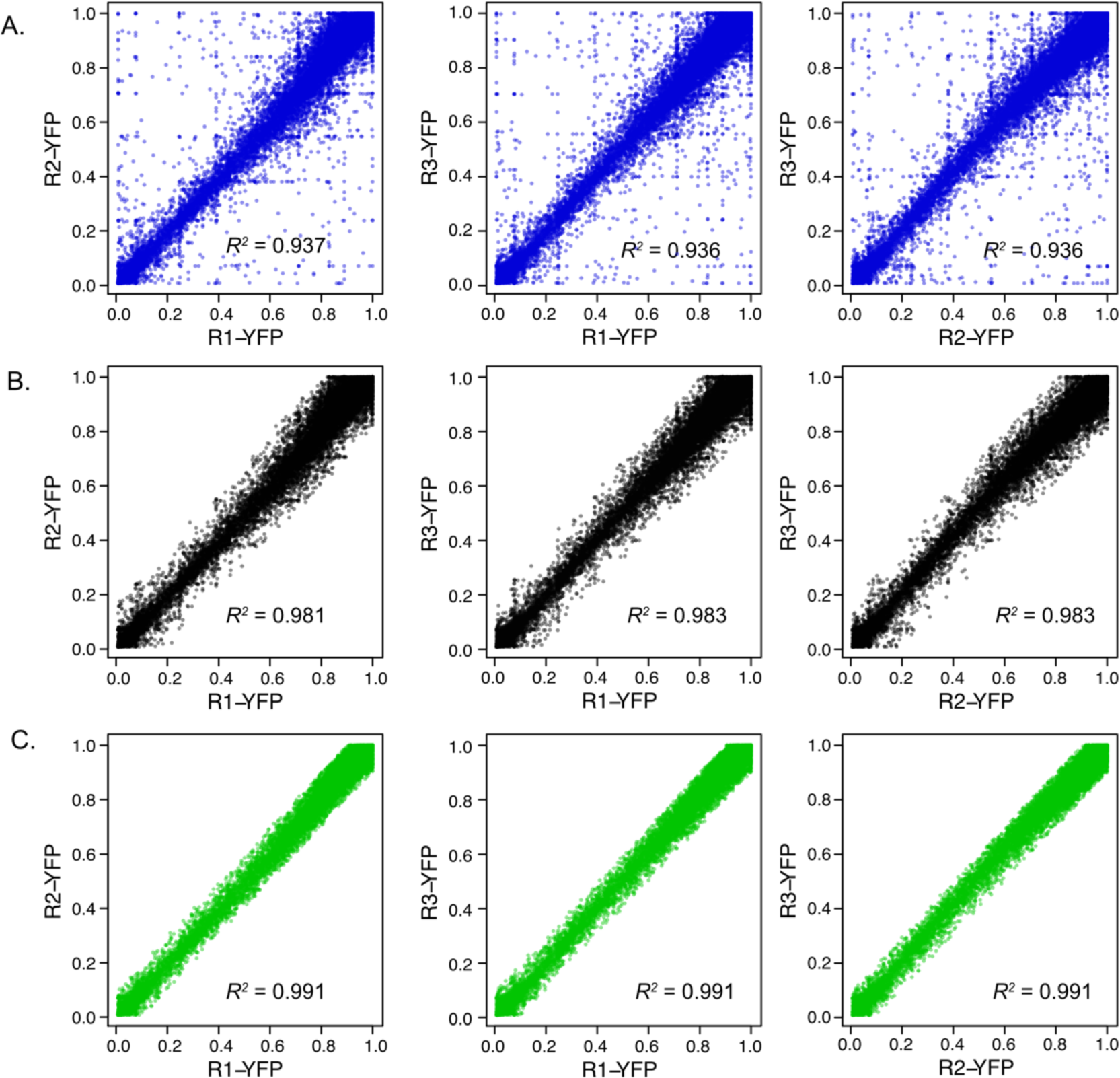
Comparison of UTR expression measurements using FACS. **A.** Raw data from three replicates are plotted against each other, with squared Pearson’s R values shown. The standard deviation across the three replicates was used to filter the data to remove UTRs with noisy measurements. Results after filtering are shown with standard deviation threshold < 0.1 (B.) and < 0.06 (C.) shown above. The 0.05 threshold was used to select UTRs with consistent YFP measurements for subsequent analysis of uORF activity using FACS-uORF (WT ∕ AAG)

**Figure S2.**
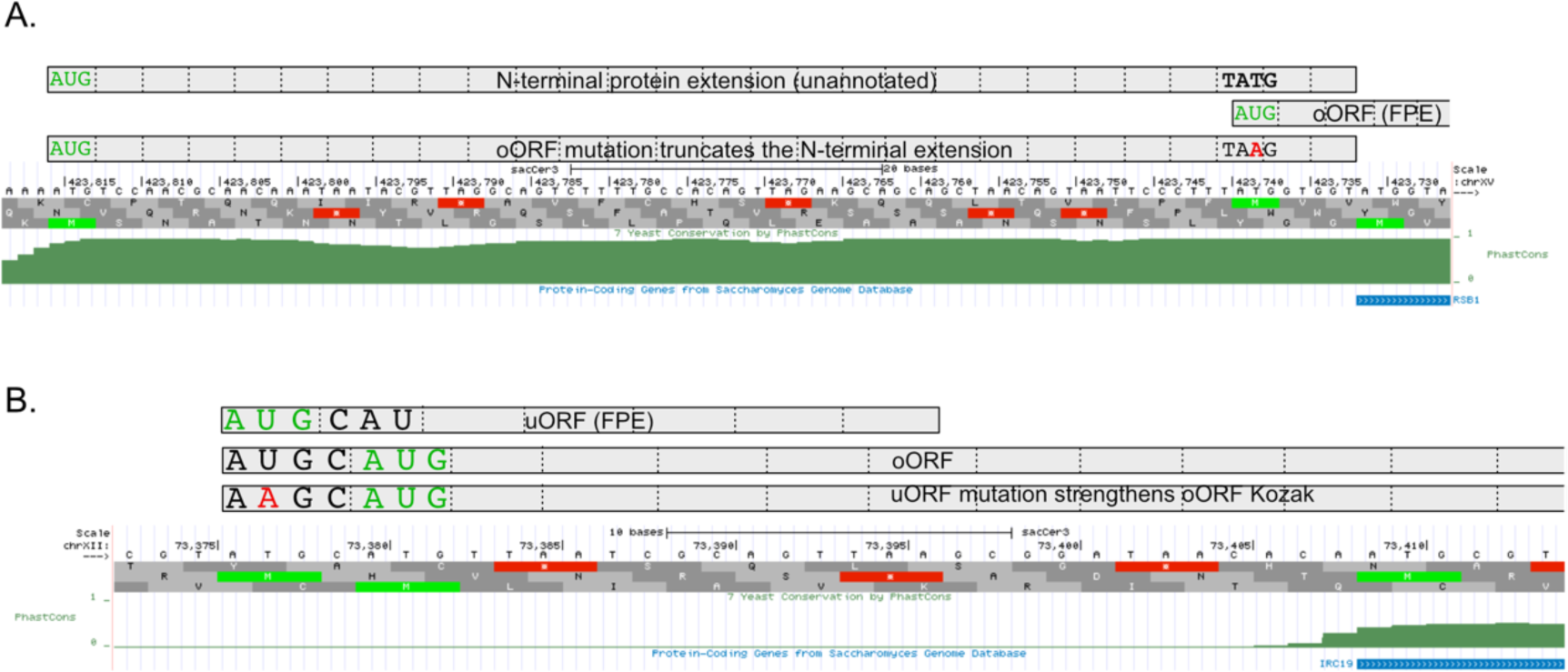
Examples of false-positive uORF enhancers. **A.** RSB1 has two upstream AUGs. The first encodes a highly conserved N-terminal protein extension that may be the true gene start codon (annotation error). The second uAUG encodes an overlapping uORF (oORF) close to the annotated gene start codon. Mutation of the oORF start codon (AUG > AAG) creates a UAA stop codon in the unannotated N-terminal extension, converting it to a uORF. This likely explains why the oORF appears to be an enhancer, as the oORF start codon mutation decreases expression. B. IRC19 also has two uAUGs. The first encodes a uORF and the second encodes an oORF. Mutation of the first uORF AUG start codon to AAG places an A in the -3 position of the oORF start codon, which strengthens the Kozak motif for the oORF. This is another common source of likely false-positive enhancer uORFs. Such cases are listed in Table S2 and were removed from all further analyses. Images created using the UCSC genome browser

**Figure S3.**
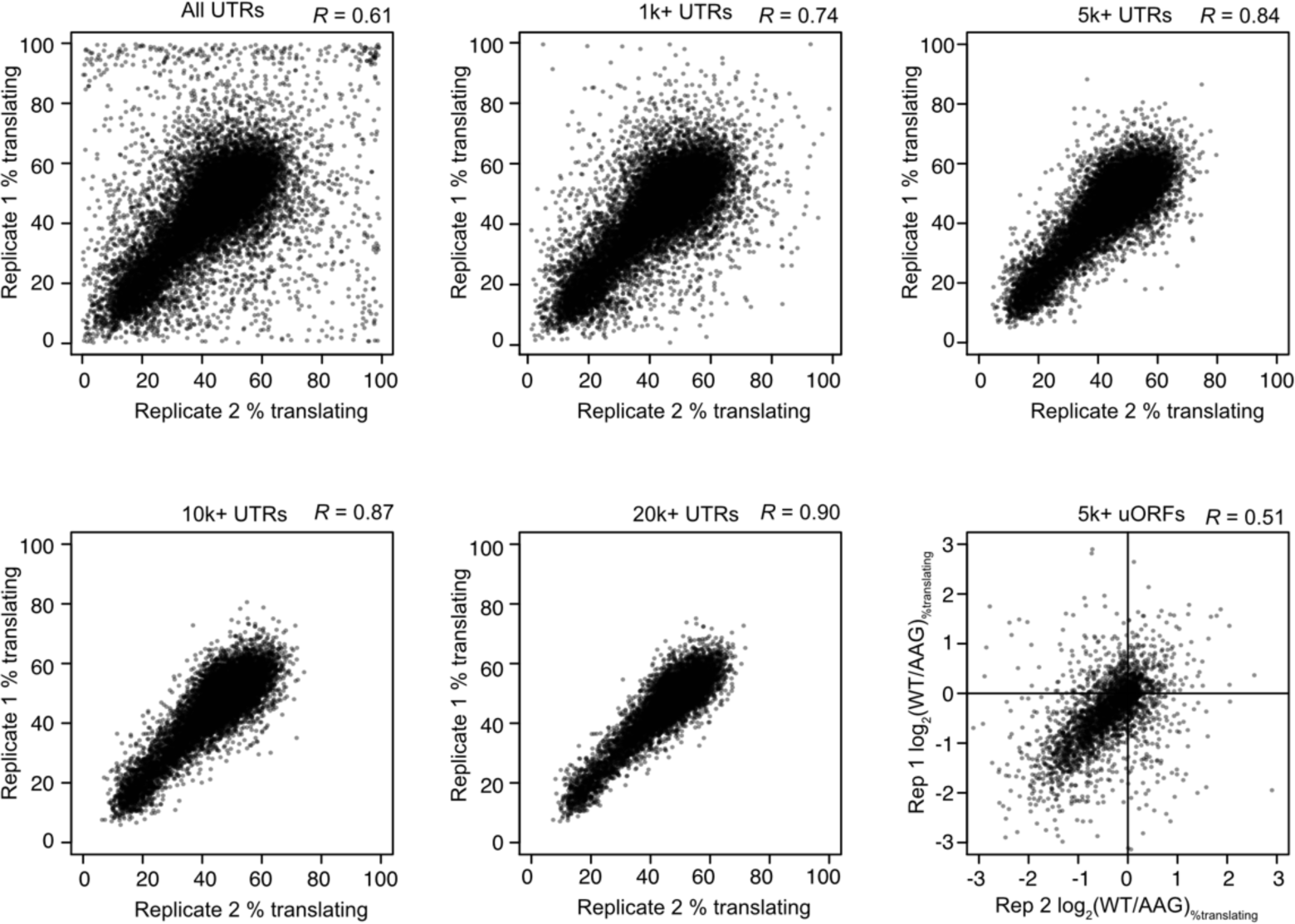
Reproducibility of PoLib-seq estimates of translation efficiency and uORF activity. Scatter plots show a comparison of two replicates of PoLib-seq data for all UTRs tested (top, left) and UTRs that exceeded read-count thresholds (1,000, 5,000, 10,000, and 20,000 total reads). Pearson’s R correlation constants are shown above each plot. The 5,000 read threshold was chosen for uORF analyses. Lower right plot shows a comparison of uORF activities as measured In the two replicates

**Figure S4.**
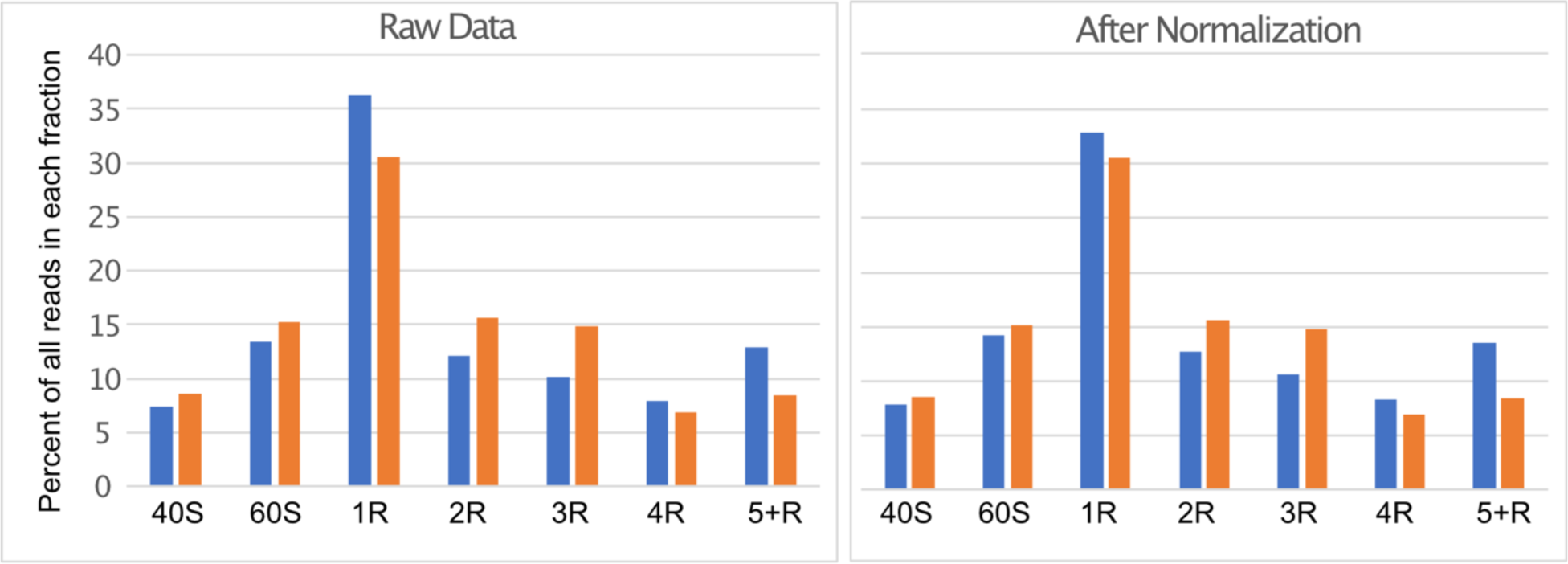
Distribution of PoLib-seq reads. Libraries were pooled based on the relative amounts of total RNA recovered from each fraction of the sucrose gradients (40S, 60S, Monosome (1 R), Disome (2R), etc.). The bar graph on the left shows the raw data from replicates 1 (blue) and two (orange). For analysis of uORF functions, reads were grouped into translating (2R, 3R, 4R, 5+R) and non-translating (40S, 60S, and 1R) fractions. Replicate 1 was normalized by downsampling the monosome fraction by a factor of 0.8626 (see methods) so that the total fraction of translating reads in each replicate was 45%, as depicted in the bar graph on the right.

## References

Amrani N, Ganesan R, Kervestin S, Mangus DA, Ghosh S, Jacobson A. 2004. A faux 3′-UTR promotes aberrant termination and triggers nonsense-mediated mRNA decay. Nature 432:112–118. doi:10.1038/nature03060

Arribere JA, Gilbert W V. 2013. Roles for transcript leaders in translation and mRNA decay revealed by transcript leader sequencing. Genome Res 23:977–987.

Barbosa C, Peixeiro I, Romão L. 2013. Gene expression regulation by upstream open reading frames and human disease. PLoS Genet 9:e1003529.

Beznosková P, Cuchalová L, Wagner S, Shoemaker CJ, Gunišová S, von der Haar T, Valášek LS. 2013. Translation Initiation Factors eIF3 and HCR1 Control Translation Termination and Stop Codon Read-Through in Yeast Cells. PLoS Genet 9:e1003962.

Bohlen J, Fenzl K, Kramer G, Bukau B, Teleman AA. 2020. Selective 40S Footprinting Reveals Cap-Tethered Ribosome Scanning in Human Cells. Mol Cell 79:561–574.e5. doi:10.1016/j.molcel.2020.06.005

Bonetti B, Fu L, Moon J, Bedwell D. 1995. The Efficiency of Translation Termination is Determined by a Synergistic Interplay Between Upstream and Downstream Sequences in Saccharomyces cerevisiae. J Mol Biol 251:334–345.

Brar GA, Yassour M, Friedman N, Regev A, Ingolia NT, Weissman JS. 2011. High-Resolution View of the Yeast Meiotic Program Revealed by Ribosome Profiling. *Sci (New York*, NY) 335:552–557.

Brown A, Shao S, Murray J, Hegde RS, Ramakrishnan V. 2015. Structural basis for stop codon recognition in eukaryotes. Nature 524:493–496. doi:10.1038/nature14896

Calvo SE, Pagliarini DJ, Mootha VK. 2009. Upstream open reading frames cause widespread reduction of protein expression and are polymorphic among humans. Proc Natl Acad Sci U S A 106:7507–7512.

Cao D, Parker R. 2003. Computational modeling and experimental analysis of nonsense-mediated decay in yeast. Cell 113:533–545. doi:10.1016/S0092-8674(03)00353-2

Celik A, Baker R, He F, Jacobson A. 2017. High-resolution profiling of NMD targets in yeast reveals translational fidelity as a basis for substrate selection. *RNA (New York*, NY) 23:735– 748.

Chew G-L, Pauli A, Schier AF. 2016. Conservation of uORF repressiveness and sequence features in mouse, human and zebrafish. Nat Commun 7:11663.

Corley M, Solem A, Phillips G, Lackey L, Ziehr B, Vincent HA, Mustoe AM, Ramos SB V, Weeks KM, Moorman NJ, Laederach A. 2017. An RNA structure-mediated, posttranscriptional model of human α-1-antitrypsin expression. Proc Natl Acad Sci U S A 114:E10244–E10253.

Cridge AG, Crowe-Mcauliffe C, Mathew SF, Tate WP. 2018. Eukaryotic translational termination efficiency is influenced by the 3′ nucleotides within the ribosomal mRNA channel. Nucleic Acids Res 46:1927–1944. doi:10.1093/nar/gkx1315

Culbertson MR, Leeds PF. 2003. Looking at mRNA decay pathways through the window of molecular evolution. Curr Opin Genet Dev 13:207–214. doi:10.1016/S0959- 437X(03)00014-5

Cuperus JT, Groves B, Kuchina A, Rosenberg AB, Jojic N, Fields S, Seelig G. 2017. Deep learning of the regulatory grammar of yeast 5’ untranslated regions from 500,000 random sequences. Genome Res 27:2015–2024.

Czaplinski K, Ruiz-Echevarria MJ, González CI, Peltz SW. 1999. Should we kill the messenger? The role of the surveillance complex in translation termination and mRNA turnover. BioEssays 21:685–696. doi:10.1002/(SICI)1521-1878(199908)21:8<685::AID-BIES8>3.0.CO;2-4

Dvir S, Velten L, Sharon E, Zeevi D, Carey LB, Weinberger A, Segal E. 2013. Deciphering the rules by which 5’-UTR sequences affect protein expression in yeast. Proc Natl Acad Sci U S A 110:E2792–801.

Eberle AB, Stalder L, Mathys H, Orozco RZ, Mühlemann O. 2008. Posttranscriptional gene regulation by spatial rearrangement of the 3′ untranslated region. PLoS Biol 6:849–859. doi:10.1371/journal.pbio.0060092

Eisenberg AR, Higdon AL, Hollerer I, Kellis M, Jovanovic M, Brar GA, Eisenberg AR, Higdon AL, Hollerer I, Fields AP, Jungreis I, Diamond PD. 2020. Article Translation Initiation Site Profiling Reveals Widespread Synthesis of Non-AUG-Initiated Protein Isoforms in Yeast Translation Initiation Site Profiling Reveals Widespread Synthesis of Non-AUG-Initiated Protein Isoforms in Yeast. Cell Syst 1–16. doi:10.1016/j.cels.2020.06.011

Ferreira JP, Noderer WL, Diaz de Arce AJ, Wang CL. 2014. Engineering ribosomal leaky scanning and upstream open reading frames for precise control of protein translation. Bioengineered 5:186–192.

Gerashchenko M V, Gladyshev VN. 2014. Translation inhibitors cause abnormalities in ribosome profiling experiments. Nucleic Acids Res 42:e134.

González CI, Ruiz-Echevarría MJ, Vasudevan S, Henry MF, Peltz SW. 2000. The yeast hnRNP-like protein Hrp1/Nab4 marks a transcript for nonsense-mediated mRNA decay. Mol Cell 5:489–499. doi:10.1016/S1097-2765(00)80443-8

Gorgoni B, Zhao YB, Krishnan J, Stansfield I. 2019. Destabilization of Eukaryote mRNAs by 5’ Proximal Stop Codons Can Occur Independently of the Nonsense-Mediated mRNA Decay Pathway. Cells 8. doi:10.3390/cells8080800

Grant CM, Miller PF, Hinnebusch AG. 2012. Requirements for intercistronic distance and level of eukaryotic initiation factor 2 activity in reinitiation on GCN4 mRNA vary with the downstream cistron. Mol Cell Biol 14:2616–2628. doi:10.1128/mcb.14.4.2616

Guenther U-P, Weinberg DE, Zubradt MM, Tedeschi FA, Stawicki BN, Zagore LL, Brar GA, Licatalosi DD, Bartel DP, Weissman JS, Jankowsky E. 2018. The helicase Ded1p controls use of near-cognate translation initiation codons in 5′ UTRs. Nature 1–20.

Gunišová S, Beznosková P, Mohammad MP, Vlčková V, Valášek LS. 2016. In-depth analysis of cis-determinants that either promote or inhibit reinitiation on GCN4mRNA after translation of its four short uORFs. *RNA (New York*, NY) 22:542–558.

Gunišová S, Hronová V, Mohammad MP, Hinnebusch AG, Valášek LS. 2017. Translation reinitiation in microbes and higher eukaryotes. FEMS Microbiol Rev.

Hastie T, Tibshirani R, Freidman J. 2009. The Elements of Statistical Learning: Data Mining, Inference, and Prediction. Springer.

Hinnebusch AG. 2005. Translational regulation of GCN4 and the general amino acid control of yeast. Annu Rev Microbiol 59:407–450.

Hinnebusch AG, Ivanov IP, Sonenberg N. 2016. Translational control by 5’-untranslated regions of eukaryotic mRNAs. *Sci (New York*, NY) 352:1413–1416.

Ingolia NT, Ghaemmaghami S, Newman JRS, Weissman JS. 2009. Genome-wide analysis in vivo of translation with nucleotide resolution using ribosome profiling. *Sci (New York*, NY) 324:218–223.

Ingolia NT, Lareau LF, Weissman JS. 2011. Ribosome Profiling of Mouse Embryonic Stem Cells Reveals the Complexity and Dynamics of Mammalian Proteomes. Cell 147:789–802.

Johnstone TG, Bazzini AA, Giraldez AJ. 2016. Upstream ORFs are prevalent translational repressors in vertebrates. EMBO J.

Kearse MG, Goldman DH, Choi J, Nwaezeapu C, Liang D, Green KM, Goldstrohm AC, Todd PK, Green R, Wilusz JE. 2019. Ribosome queuing enables non-AUG translation to be resistant to multiple protein synthesis inhibitors. Genes Dev 33:871–885. doi:10.1101/gad.324715.119

Kearse MG, Wilusz JE. 2017. Non-AUG translation: a new start for protein synthesis in eukaryotes. Genes Dev 31:1717–1731.

Keeling KM, Lanier J, Du M, Salas-Marco J, Gao L, Kaenjak-Angeletti A, Bedwell DM. 2004. Leaky termination at premature stop codons antagonizes nonsense-mediated mRNA decay in S. cerevisiae. Rna 10:691–703. doi:10.1261/rna.5147804

Kolitz SE, Takacs JE, Lorsch JR. 2009. Kinetic and thermodynamic analysis of the role of start codon/anticodon base pairing during eukaryotic translation initiation. *RNA (New York*, NY) 15:138–152.

Kozak M. 1991. Structural features in eukaryotic mRNAs that modulate the initiation of translation. J Biol Chem 266:19867–19870.

Lareau LF, Hite DH, Hogan GJ, Brown PO. 2014. Distinct stages of the translation elongation cycle revealed by sequencing ribosome-protected mRNA fragments. Elife 3:e01257.

Lee DSM, Park J, Kromer A, Rader DJ, Ritchie MD, Ghanem LR, Barash Y. 2020. Disrupting upstream translation in mRNAs leads to loss-of-function associated with human disease. bioRxiv. doi:10.1101/2020.09.09.287912

Letzring DP, Wolf AS, Brule CE, Grayhack EJ. 2013. Translation of CGA codon repeats in yeast involves quality control components and ribosomal protein L1. *RNA (New York*, NY) 19:1208–1217.

Li JJ, Chew GL, Biggin MD. 2019. Quantitative principles of cis-translational control by general mRNA sequence features in eukaryotes. Genome Biol 20:1–24. doi:10.1186/s13059-019-1761-9

Lin Y, May GE, Kready H, Nazzaro L, Mao M, Spealman P, Creeger Y, McManus CJ. 2019. Impacts of uORF codon identity and position on translation regulation. Nucleic Acids Res 2:1–10. doi:10.1093/nar/gkz681

Lindeboom RGH, Supek F, Lehner B. 2016. The rules and impact of nonsense-mediated mRNA decay in human cancers. Nat Genet 48:1112–1118. doi:10.1038/ng.3664

Lu Z, Lin Z. 2019. Pervasive and dynamic transcription initiation in Saccharomyces cerevisiae. Genome Res 29:1198–1210. doi:10.1101/gr.245456.118

Magoč T, Salzberg SL. 2011. FLASH: Fast length adjustment of short reads to improve genome assemblies. Bioinformatics 27:2957–2963. doi:10.1093/bioinformatics/btr507

Martin M. 2011. Cutadapt removes adapter sequences from high-throughput sequencing reads. EMBnet.journal 17:10–12.

McGillivray P, Ault R, Pawashe M, Kitchen R, Balasubramanian S, Gerstein M. 2018. A comprehensive catalog of predicted functional upstream open reading frames in humans. Nucleic Acids Res 46:3326–3338.

McManus CJ, May GE, Spealman P, Shteyman A. 2014. Ribosome profiling reveals post-transcriptional buffering of divergent gene expression in yeast. Genome Res 24:422–430.

Meaux S, van Hoof A, Baker KE. 2008. Nonsense-Mediated mRNA Decay in Yeast Does Not Require PAB1 or a Poly(A) Tail. Mol Cell 29:134–140. doi:10.1016/j.molcel.2007.10.031

Mueller PP, Hinnebusch AG. 1986. Multiple upstream AUG codons mediate translational control of GCN4. Cell 45:201–207. doi:10.1016/0092-8674(86)90384-3

Muhlrad D, Parker R. 1999. Aberrant mRNAs with extended 3’ UTRs are substrates for rapid degradation by mRNA surveillance. Rna 5:1299–1307. doi:10.1017/S1355838299990829

Mustoe AM, Corley M, Laederach A, Weeks KM. 2018. Messenger RNA Structure Regulates Translation Initiation: A Mechanism Exploited from Bacteria to Humans. Biochemistry 57:3537–3539. doi:10.1021/acs.biochem.8b00395

Namy O, Hatin I, Rousset JP. 2001. Impact of the six nucleotides downstream of the stop codon on translation termination. EMBO Rep 2:787–793. doi:10.1093/embo-reports/kve176

Noderer WL, Flockhart RJ, Bhaduri A, Diaz de Arce AJ, Zhang J, Khavari PA, Wang CL. 2014. Quantitative analysis of mammalian translation initiation sites by FACS-seq. Mol Syst Biol 10:748.

Oertlin C, Lorent J, Gandin V, Murie C, Masvidal L, Cargnello M, Furic L, Topisirovic I, Larsson O. 2017. Genome-wide analysis of differential translation and differential translational buffering using anota2seq. *bioRxiv*.

Pelechano V, Wei W, Steinmetz LM. 2013. Extensive transcriptional heterogeneity revealed by isoform profiling. Nature 1–7.

Peltz SW, Brown AH, Jacobson A. 1993. mRNA destabilization triggered by premature translational termination depends on at least three cis-acting sequence elements and one trans-acting factor. Genes Dev 7:1737–1754. doi:10.1101/gad.7.9.1737

Raimondeau E, Bufton JC, Schaffitzel C. 2018. New insights into the interplay between the translation machinery and nonsense-mediated mRNA decay factors. Biochem Soc Trans 46:503–512. doi:10.1042/BST20170427

Rojas-Duran MF, Gilbert W V. 2012. Alternative transcription start site selection leads to large differences in translation activity in yeast. *RNA (New York*, NY) 18:2299–2305.

Ruiz-Echevarría MJ, González CI, Peltz SW. 1998. Identifying the right stop: Determining how the surveillance complex recognizes and degrades an aberrant mRNA. EMBO J 17:575– 589. doi:10.1093/emboj/17.2.575

Ruiz-Echevarría MJ, Peltz SW. 2000. The RNA binding protein Pub1 modulates the stability of transcripts containing upstream open reading frames. Cell 101:741–751.

Ryabova LA, Hohn T. 2000. Ribosome shunting in the cauliflower mosaic virus 35S RNA leader is a special case of reinitiation of translation functioning in plant and animal systems. Genes Dev 14:817–829.

Sample PJ, Wang B, Reid DW, Presnyak V, McFadyen IJ, Morris DR, Seelig G. 2019. Human 5′ UTR design and variant effect prediction from a massively parallel translation assay. Nat Biotechnol 37:803–809. doi:10.1038/s41587-019-0164-5

Scannell DR, Zill OA, Rokas A, Payen C, Dunham MJ, Eisen MB, Rine J, Johnston M, Hittinger CT. 2011. The awesome power of yeast evolutionary genetics: new genome sequences and strain resources for the Saccharomyces sensu stricto genus. G3 Genes, Genomes, Genet 1:11–25.

Sen ND, Gupta N, K Archer S, Preiss T, Lorsch JR, Hinnebusch AG. 2019. Functional interplay between DEAD-box RNA helicases Ded1 and Dbp1 in preinitiation complex attachment and scanning on structured mRNAs in vivo. Nucleic Acids Res 47:8785–8806. doi:10.1093/nar/gkz595

Sen ND, Zhou F, Ingolia NT, Hinnebusch AG. 2015. Genome-wide analysis of translational efficiency reveals distinct but overlapping functions of yeast DEAD-box RNA helicases Ded1 and eIF4A. Genome Res.

Sharma D, Jankowsky E. 2014. The Ded1/DDX3 subfamily of DEAD-box RNA helicases. Crit Rev Biochem Mol Biol 49:343–360. doi:10.3109/10409238.2014.931339

Smith JE, Alvarez-Dominguez JR, Kline N, Huynh NJ, Geisler S, Hu W, Coller J, Baker KE. 2014. Translation of Small Open Reading Frames within Unannotated RNA Transcripts in Saccharomyces cerevisiae. Cell Reports 7:1858–1866.

Spealman P, Naik AW, May GE, Kuersten S, Freeberg L, Murphy RF, McManus J. 2018. Conserved non-AUG uORFs revealed by a novel regression analysis of ribosome profiling data. Genome Res 28:214–222. doi:10.1101/gr.221507.117

Spealman P, Naik AW, May GE, Kuersten S, Freeberg L, Murphy RF, McManus J. 2017. Conserved non-AUG uORFs revealed by a novel regression analysis of ribosome profiling data. Genome Res.

Takacs JE, Neary TB, Ingolia NT, Saini AK, Martin-Marcos P, Pelletier J, Hinnebusch AG, Lorsch JR. 2011. Identification of compounds that decrease the fidelity of start codon recognition by the eukaryotic translational machinery. *RNA (New York*, NY) 17:439–452.

Tang H-L, Yeh L-S, Chen N-K, Ripmaster T, Schimmel P, Wang C-C. 2004. Translation of a yeast mitochondrial tRNA synthetase initiated at redundant non-AUG codons. J Biol Chem 279:49656–49663.

Vattem KM, Wek RC. 2004. Reinitiation involving upstream ORFs regulates ATF4 mRNA translation in mammalian cells. Proc Natl Acad Sci 101:11269–11274. doi:10.1073/pnas.0400541101

Wagner S, Herrmannová A, Hronová V, Gunišová S, Sen ND, Hannan RD, Hinnebusch AG, Shirokikh NE, Preiss T, Valášek LS. 2020. Selective Translation Complex Profiling Reveals Staged Initiation and Co-translational Assembly of Initiation Factor Complexes. Mol Cell 79:546–560.e7. doi:10.1016/j.molcel.2020.06.004

Wethmar K. 2014. The regulatory potential of upstream open reading frames in eukaryotic gene expression. Wiley Interdiscip Rev RNA 5:765–778.

Wethmar K, Barbosa-Silva A, Andrade-Navarro MA, Leutz A. 2014. uORFdb--a comprehensive literature database on eukaryotic uORF biology. Nucleic Acids Res 42:D60–7.

Wethmar K, Smink JJ, Leutz A. 2010. Upstream open reading frames: molecular switches in (patho)physiology. Bioessays 32:885–893.

Young SK, Wek RC. 2016. Upstream Open Reading Frames Differentially Regulate Gene-specific Translation in the Integrated Stress Response. J Biol Chem 291:16927–16935.

Zhang Fan, Hinnebusch AG. 2011. An upstream ORF with non-AUG start codon is translated in vivo but dispensable for translational control of GCN4 mRNA. Nucleic Acids Res 39:3128– 3140. doi:10.1093/nar/gkq1251

Zhang F, Hinnebusch AG. 2011. An upstream ORF with non-AUG start codon is translated in vivo but dispensable for translational control of GCN4 mRNA. Nucleic Acids Res 39:3128– 3140.

Zhang H, Dou S, He F, Luo J, Wei L, Lu J. 2018. Genome-wide maps of ribosomal occupancy provide insights into adaptive evolution and regulatory roles of uORFs during Drosophila development. PLoS Biol 16:e2003903.

Zhang H, Wang Y, Lu J. 2019. Function and Evolution of Upstream ORFs in Eukaryotes. Trends Biochem Sci 44:782–794. doi:10.1016/j.tibs.2019.03.002

Zhang J, Maquat LE. 1997. Evidence that translation reinitiation abrogates nonsense-mediated mRNA decay in mammalian cells. EMBO J 16:826–833. doi:10.1093/emboj/16.4.826

Zhang Z, Dietrich FS. 2005a. Identification and characterization of upstream open reading frames (uORF) in the 5’ untranslated regions (UTR) of genes in Saccharomyces cerevisiae. Curr Genet 48:77–87.

Zhang Z, Dietrich FS. 2005b. Mapping of transcription start sites in Saccharomyces cerevisiae using 5’ SAGE. Nucleic Acids Res 33:2838–2851.

Zydowicz-Machtel P, Swiatkowska A, Popenda L, Gorska A, Ciesiołka J. 2018. Variants of the 5′-terminal region of p53 mRNA influence the ribosomal scanning and translation efficiency. Sci Rep 8:1–14. doi:10.1038/s41598-018-20010-2

